# Anti-amyloid beta therapy resolves stroke recovery impairment caused by Alzheimer’s disease

**DOI:** 10.1101/2025.05.12.653388

**Authors:** Seiichiro Sakai, Kento Otani, Jun Tsuyama, Yuichiro Hara, Ko Abe, Teppei Shimamura, Ito Kawakami, Takashi Komori, Hideya Kawaji, Koji Hase, Takashi Saito, Takashi Shichita

**Affiliations:** Department of Neuroinflammation and Repair, Medical Research Laboratory, Institute of Science Tokyo, Tokyo 113-8510, Japan and Core Research for Evolutionary Medical Science and Technology (CREST), Japan Agency for Medical Research and Development (AMED), Tokyo 100-0004, Japan; Stroke Renaissance Project, Tokyo Metropolitan Institute of Medical Science, Tokyo 156-8506, Japan; Division of Biochemistry, Faculty of Pharmacy and Graduate School of Pharmaceutical Science, Keio University, Tokyo 105-8512, Japan; Research Center for Genome & Medical Sciences, Tokyo Metropolitan Institute of Medical Science, Tokyo 156-8506, Japan; Department of Computational and Sytems Biology, Medical Research Laboratory, Institute of Science Tokyo, Tokyo 113-8510, Japan; Laboratory for Neuropathology, Tokyo Metropolitan Institute of Medical Science, Tokyo 156-8506, Japan; Department of Laboratory Medicine and Pathology (Neuropathology), Tokyo Metropolitan Neurological Hospital, Tokyo Metropolitan Hospital Organization, Tokyo 183-0042, Japan; Institute of Fermentation Sciences (IFeS), Faculty of Food and Agricultural Sciences, Fukushima University, Fukushima 960-1296, Japan; International Vaccine Design Center, The Institute of Medical Science, The University of Tokyo (IMSUT), Tokyo 108-8639, Japan; Department of Neurocognitive Science, Institute of Brain Science, Nagoya City University, Graduate School of Medical Sciences, Nagoya 467-8601, Japan; Department of Neuroscience and Pathobiology, Research Institute of Environmental Medicine, Nagoya University, Nagoya 464-8601, Japan; Department of Medicine and Clinical Science, Graduate School of Medical Sciences, Kyushu University, Fukuoka 812-8582, Japan

## Abstract

Stroke and dementia are common comorbidities and a growing concern causing disability in aging societies worldwide. Although anti-amyloid beta (anti-Aβ) antibodies have recently been anticipated to relieve preclinical Alzheimer’s disease pathology, we discovered that post-stroke administration of anti-Aβ antibodies restored neural repair for stroke recovery impeded by cerebral Aβ accumulation. Neuronal recovery-associated gene expression for stroke recovery was considerably impaired even by slight Aβ accumulation in murine and human brain. Slight Aβ accumulation had less impact on neurons without stroke but caused a unique myeloid immunity after an ischemic stroke that enhanced the inflammatory cascades impeding neural repair for stroke recovery. Aducanumab administration after ischemic stroke prevented formation of this malignant myeloid immunity, resolving the impairment of post-stroke neural repair caused by cerebral Aβ accumulation. Thus, our study has revealed the ability of anti-Aβ therapies to restore functional recovery after a stroke with cerebral Aβ accumulation.

## Introduction

As a result of an aging society worldwide, the incidence of dementia and stroke is increasing, and dementia and stroke are major causes of disability and increasing medical expenditure^1–3^. It is now more common to detect both dementia and stroke pathologies in the brains of older people^4^. Although approximately sixty percent of neurocognitive disorders are due to Alzheimer’s disease, many studies have reported that 40 to 50 % of patients with Alzheimer’s disease also have cerebrovascular lesions or evidence of a previous stroke in their brain tissue^4–7^. Stroke is recognized as a major factor in the progression of Alzheimer’s disease and neurocognitive deficits^8^. Thus, the relationship between stroke and Alzheimer’s disease is complicated; each progressively worsens the pathology of the other^4,8^. This relationship can render stroke therapeutics ineffective and makes it difficult to develop treatments that improve the functional prognosis of stroke patients with Alzheimer’s disease.

Recently, anti-amyloid beta (anti-Aβ) antibodies such as aducanumab and lecanemab that deplete aggregated Aβ species (oligomers, protofibrils, and insoluble fibrils) have been expected to prevent the initiation or progression of Alzheimer’s disease pathologies^9,10^. Cerebral accumulation of aggregated Aβ is considered a first sign of Alzheimer’s disease pathology. Aβ positron emission tomography (Aβ PET) is a promising technique for detecting healthy elderly individuals with cerebral Aβ accumulation who are likely to develop neurocognitive deficits^11,12^. It is possible that these emerging medical technologies could be applied to enhance functional recovery after brain injuries, indicating the need for research to clarify the effect of Aβ pathologies on the molecular mechanisms underlying stroke recovery.

Cellular and molecular mechanisms underlying stroke recovery have recently been elucidated, attracting attention to developing therapeutics that promote recovery from neurological deficits and prolong healthy life expectancy^13,14^. We have previously demonstrated the drastically increased gene expression associated with recovery processes that occur after brain injury, including nervous tissue reconstruction, synaptic organization, and remyelination, in peri-infarct neurons, which are necessary for long-term neurological recovery after ischemic stroke^15^. Several reports demonstrate poor synaptic plasticity in patients with severe neurocognitive deficits, possibly explaining why stroke recovery is irreversibly impaired by advanced dementia pathologies^16^. However, it remains to be clarified whether or how a small accumulation of aggregated Aβ in brain tissue affects the cellular or molecular mechanisms underlying stroke recovery.

For this purpose, we used *App^NL-G-F^* knockin (KI) mice bearing humanized Aβ sequence with the Arctic, Swedish, and Beyreuther/Iberian mutations observed in familial Alzheimer’s disease. *App^NL-G-F^* KI mice are suitable for investigating the direct effect of cerebral Aβ accumulation on stroke recovery because reproducible cerebral Aβ accumulation is observed in an age-dependent manner. In research conducted to date, *App* transgenic mice overexpressing amyloid precursor protein (APP) have been commonly used; however, they also generate APP intracellular domain (AICD) or C-terminal fragment β (CTF-β) that may have unpredicted effects for brain pathologies^17,18^, preventing evaluation of stroke recovery with cerebral Aβ accumulation. We discovered that even a slight accumulation of Aβ in the brain tissue of *App^NL-G-F^* KI mice impaired stroke recovery. Slight cerebral Aβ accumulation did not affect neuronal function without stroke but induced the unique myeloid immunity formation after ischemic stroke, which impaired neural repair necessary for stroke recovery.

## Results

### Neuronal recovery mechanisms were impaired after ischemic stroke in *App^NL-G-F^* KI mice

First, we confirmed the deposition of aggregated Aβ in the brain tissue of *App^NL-G-F^* KI mice before induction of ischemic stroke by immunohistochemistry using mouse chimeric analogs of aducanumab (mAducanumab) specific for Aβ oligomers and fibrils^19^. The deposition of aggregated Aβ in the cerebral cortex was slight in heterozygous 2-month-old *App^NL-G-F^* KI mice but remarkably increased in 2- or 6-month-old homozygous *App^NL-G-F^* KI mice (**Fig.1a**). Elevated levels of human Aβ_42_ were also confirmed in the brain tissues of *App^NL-G-F^* KI mice (**Supplementary Fig.1a**).

**Figure 1.**
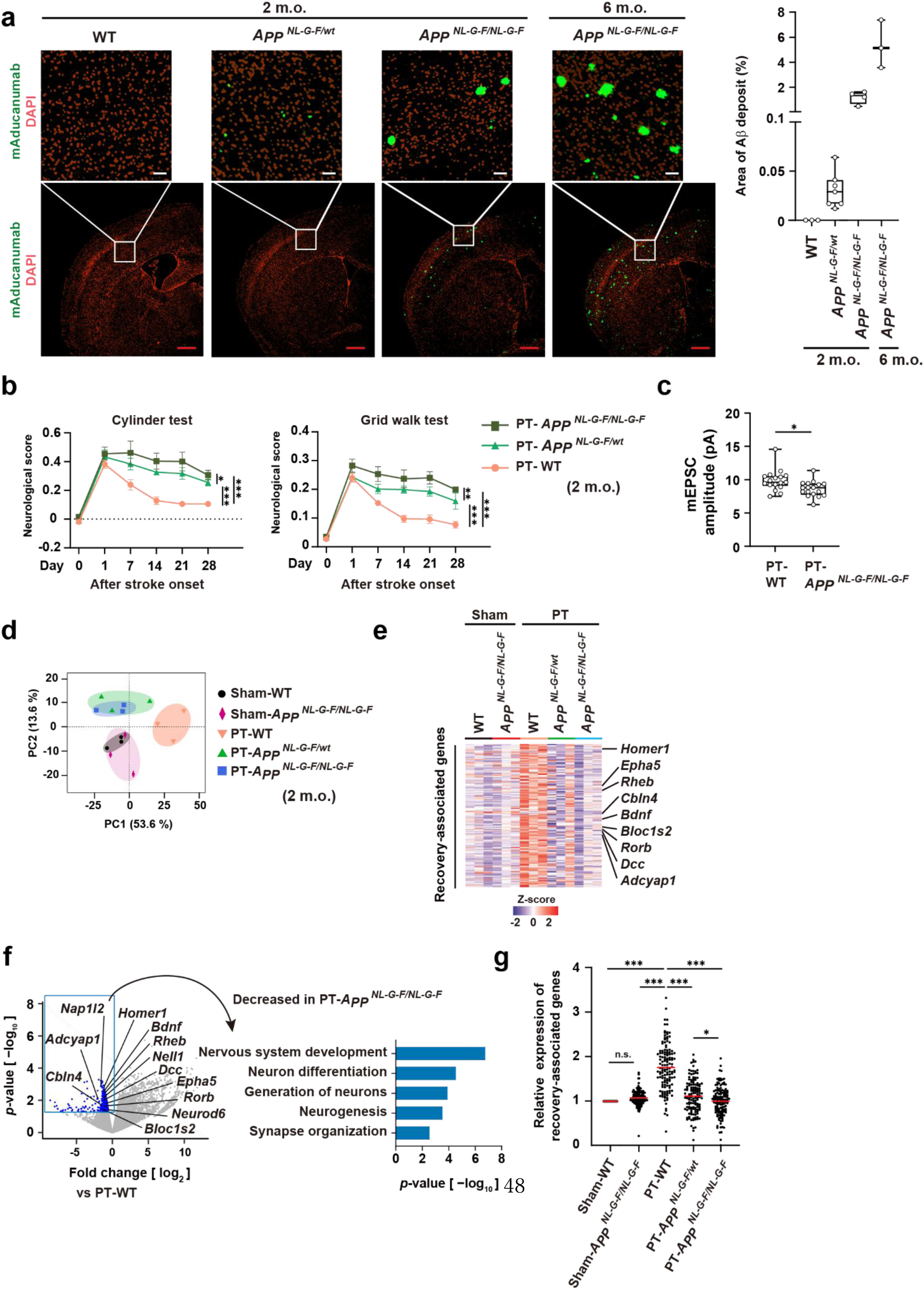
Cerebral Aβ accumulation impaired stroke recovery. (**a**) Immunohistochemistry using mAducanumab in the brain tissue before the induction of ischemic stroke (white bar: 50 μm, red bar: 500 μm, m.o.: months old). The right panel shows the ratio of Aβ-positive area. (**b**) The time-dependent changes of neurological deficits after ischemic stroke induced by photothrombosis (PT) in 2-month-old mice (n = 7 for PT-*App^NL-G-F/NL-G-F^*, n = 8 for PT-*App^NL-G-F/wt^*, n = 9 for PT-WT). (**c**) The amplitude of the miniature excitatory post-synaptic current (mEPSC) in the peri-infarct neurons of post-ischemic brain tissue 4 weeks after stroke onset. (**d**) PCA and (**e**) heatmaps of recovery-associated gene expression levels in neurons analyzed by bulk RNA-seq (n = 3 for each group). Peri-infarct neurons of PT-mice were isolated 7 days after stroke onset (**d**,**e**). Recovery-associated genes were defined as genes included in the gene ontology terms “Nervous system reconstruction,” “Synaptic organization,” and “Remyelination”^15^, whose expression levels were significantly increased in post-ischemic day 7 neurons of PT-WT mice compared to the neurons of sham-operated (Sham-) WT mice (**e**). (**f**) RNA-seq results of post-ischemic day 7 neurons in PT-*App^NL-G-F/NL-G-F^* mice compared to PT-WT mice. The right panel shows the gene ontology analysis of the genes whose expression levels significantly decreased in PT-*App^NL-G-F/NL-G-F^* mice compared to PT-WT mice. (**g**) Relative expression of each recovery-associated gene compared to Sham-WT. \**p* < 0.05, \*\**p* < 0.01, \*\*\**p* < 0.001 vs. PT-WT (**c**) (two-way ANOVA with Tukey’s test [**b**], two-sided Student’s *t*-test [**c**], one-way ANOVA with Tukey’s test [**g**]). Error bars represent the mean ± standard error of the mean (SEM).

No neurological deficit was observed in sham-operated 2-month-old *App^NL-G-F^* KI mice (**Supplementary Fig.1b**); however, recovery from neurological deficits after ischemic stroke induced by photothrombosis (PT) of cortical microvessels was significantly impaired compared with that of wild type (WT) mice (**Fig.1b**). There was no significant difference in infarct volume, physiological data, or survival rate between WT and *App^NL-G-F^* KI mice (**Supplementary Fig.1c**,**d** and **Supplementary Table 1**). Impaired post-stroke recovery was also observed in 6-month-old *App^NL-G-F^* KI mice (**Supplementary Fig.1e**), although trivial neurocognitive deficits were reported in 6-month-old *App^NL-G-F^* KI mice without stroke but not in 2-month-old *App^NL-G-F^* KI mice^20–22^. These results indicated that even a slight deposition of aggregated Aβ in the brain tissue impaired stroke recovery.

Because post-stroke synaptic organization was dysfunctional in the peri-infarct neurons of 2-month-old *App^NL-G-F^* KI mice 4 weeks after stroke onset (**Fig.1c**), we investigated the gene expression profiles of peri-infarct neurons isolated from ischemic brain on day 7 after stroke onset by bulk RNA-seq analysis. As we reported previously^15^, ischemic stroke induced gene expression associated with stroke recovery processes involving nervous tissue reconstruction, synaptic plasticity, and remyelination in peri-infarct neurons. Principal component analysis (PCA) of recovery-associated gene expression showed that a small amount of Aβ accumulation in 2-month-old homozygous *App^NL-G-F^* KI mice had little impact in the neurons of the sham-operated brain (**Fig.1d**). In contrast, ischemic stroke strongly influenced recovery-associated gene expression in peri-infarct neurons, and these differed between WT and *App^NL-G-F^* KI mice (**Fig.1d**).

Expression of these recovery-associated genes after ischemic stroke was impaired in both heterozygous and homozygous *App^NL-G-F^* KI mice compared with WT mice (**Fig.1e**). Indeed, gene ontologies associated with the recovery processes, such as nervous tissue development and synapse organization, were enriched in genes whose expression was significantly decreased in *App^NL-G-F^* KI mice with ischemic stroke, compared with WT mice with ischemic stroke (**Fig.1f**). As shown in **Fig.1g**, slight cerebral Aβ accumulation considerably decreased neuronal recovery-associated gene expression, and homozygous rather than heterozygous *App^NL-G-F^* KI mice showed severe impairment of recovery-associated gene expression, revealing that cerebral Aβ accumulation levels correlated with impairment of recovery-associated gene expression (**Fig.1a**,**g**). These results demonstrated that neuronal recovery mechanisms after ischemic stroke were impaired, even in mice with slight cerebral Aβ accumulation.

### Neuronal recovery-associated gene expression was impaired in human ischemic brain with Aβ accumulation

To confirm our findings in human ischemic brain tissue, ischemic stroke patients were separated into three groups: negligible, slight, or remarkable accumulation of Aβ in their brain tissue (**Fig.2a**,**b**), and we examined the immunohistochemistry of BDNF and NELL1, recovery-associated genes whose expression was impaired by cerebral Aβ accumulation in mice (**Fig.1f**). Compared to the brain samples without ischemic stroke, BDNF and NELL1 expression was significantly increased in the peri-infarct neurons of ischemic stroke patients with negligible Aβ deposits (**Fig.2c−f**). However, significantly decreased BDNF and NELL1 expression was observed in stroke patients with slight or remarkable Aβ deposition, compared to those with negligible Aβ deposits (**Fig.2c−f**). Neuronal BDNF and NELL1 expression after ischemic stroke was inversely proportional to cerebral Aβ accumulation level (**Fig.2g**). Thus, gene expression associated with stroke recovery was impaired in the human brain with slight Aβ accumulation.

**Figure 2.**
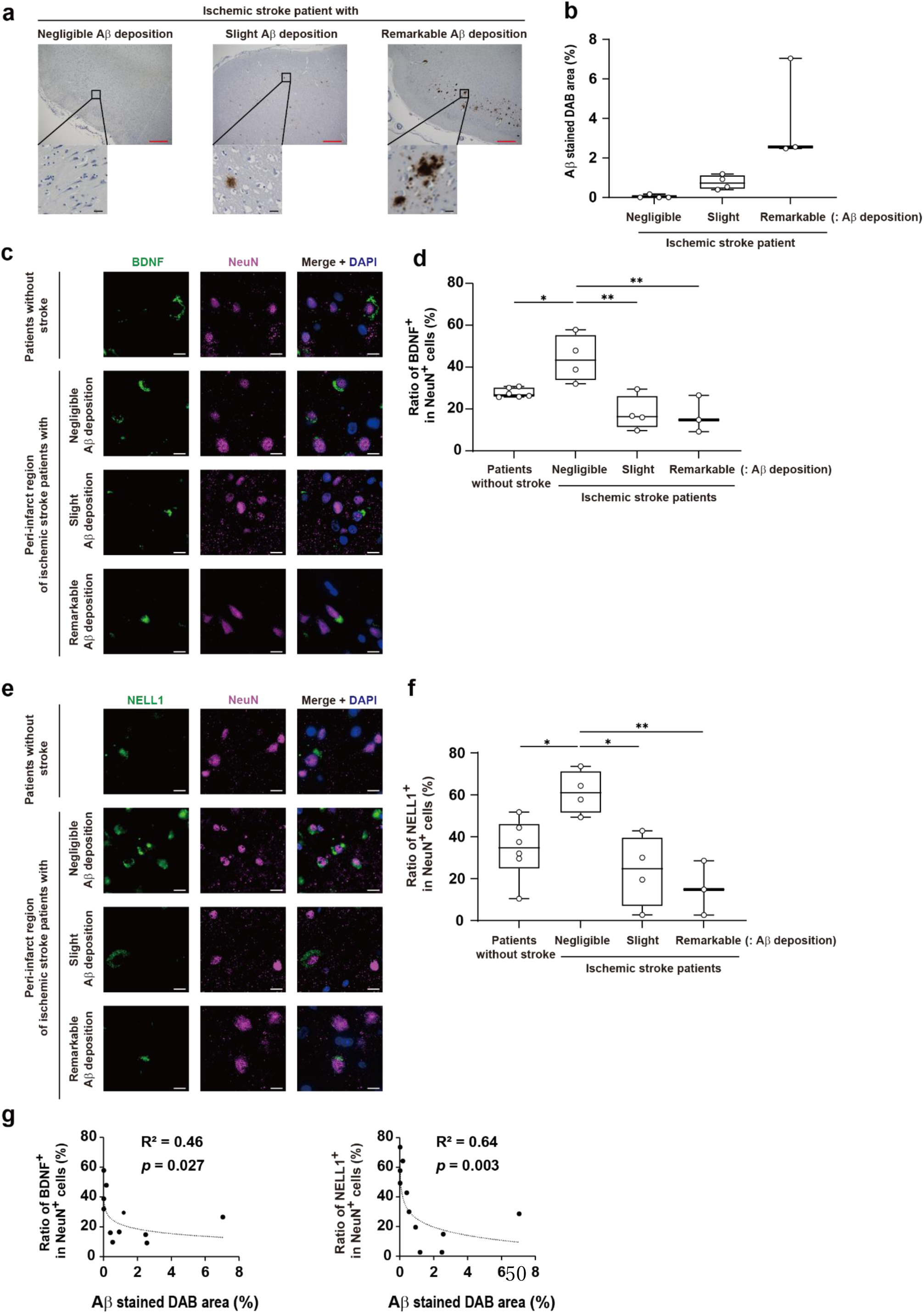
Impaired expression of neuronal recovery-associated genes in the human post-ischemic brain with Aβ accumulation. (**a**) Immunohistochemistry of Aβ in the hippocampus (red bar: 500 μm, black bar: 30 μm) and (**b**) the ratio of Aβ-positive area in the peri-infarct region and hippocampus of the human post-ischemic brain. Ischemic stroke patients were separated into three groups: negligible (< 0.2 %), slight (< 2 %), and remarkable (> 2 % of 1.5 mm^2^ area). (**c−f**) Immunohistochemistry of BDNF^+^ or NELL1^+^ neurons (bar: 10 μm) (**c**,**e**) and the ratio of BDNF^+^ or NELL1^+^ neuronal cells (**d**,**f**) in the human brain tissue. (**g**) Correlation between BDNF^+^ or NELL1^+^ neurons and the Aβ-positive area ratio. The coefficient of determination (R-squared) was calculated using regression analysis. \**p* < 0.05, \*\**p* < 0.01 (one-way ANOVA with Tukey’s test [**d**,**f**], Spearman’s rank correlation [**g**]). Error bars represent the mean ± standard error of the mean (SEM).

### Unique myeloid immunity was induced in ischemic brain with Aβ accumulation

To identify the mechanisms underlying impaired stroke recovery in 2-month-old *App^NL-G-F^* KI mice, we investigated gene expression profiles in various brain cells using single-cell RNA-seq (scRNA-seq) (**Fig.3a**). We found that the most significant number of differentially expressed genes (DEGs) were detected in myeloid cells, followed by excitatory neurons and oligodendrocytes when comparing the different brain conditions (**Fig.3b**). Gene expression associated with nervous tissue reconstruction, synaptic plasticity, and remyelination was remarkably increased in excitatory neurons and oligodendrocytes 7 days after ischemic stroke onset but this was not observed in the post-ischemic brain of *App^NL-G-F^* KI mice (**Fig.3c**,**d**).

**Figure 3.**
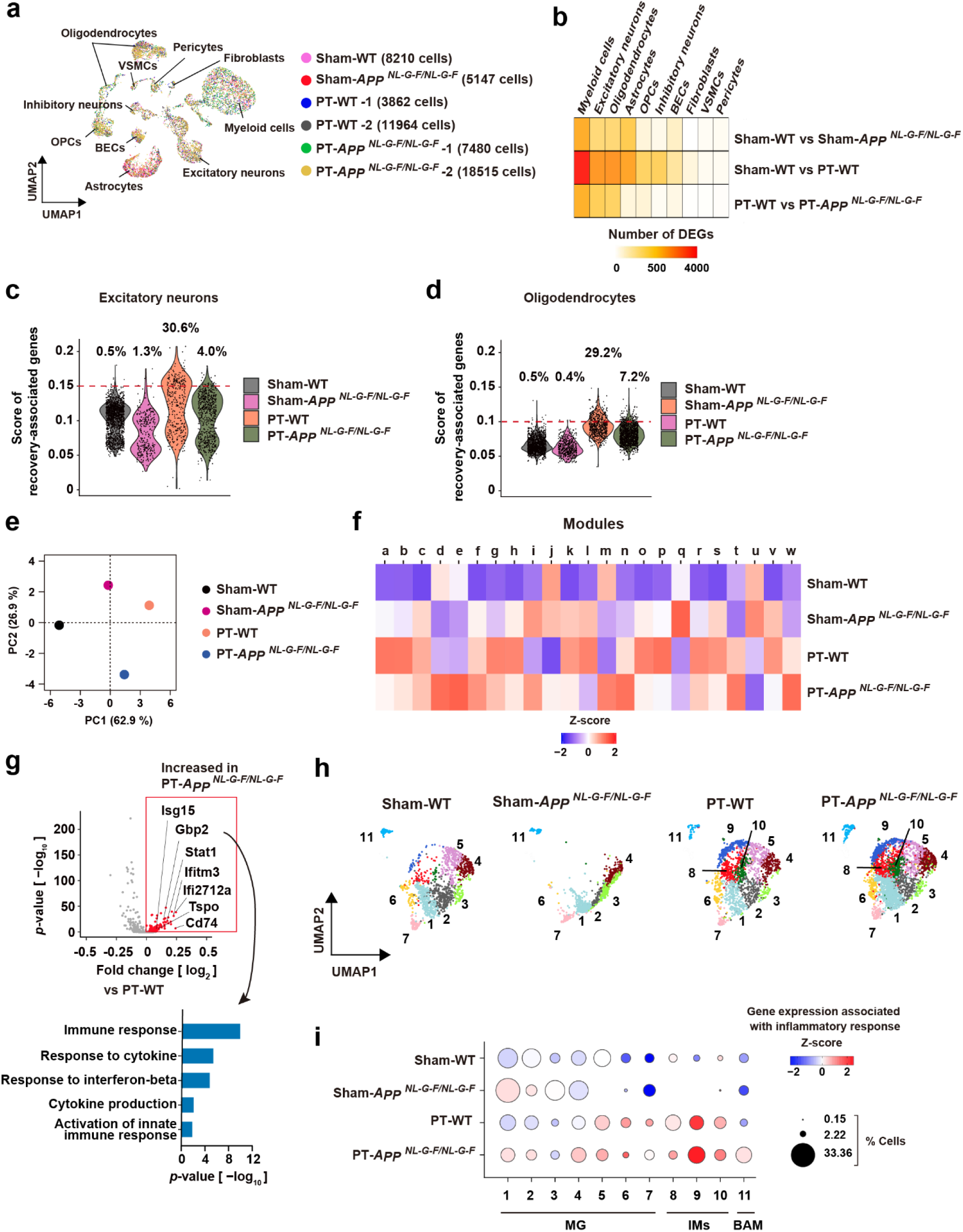
Cerebral Aβ accumulation induced the unique pro-inflammatory myeloid immunity after ischemic stroke. (**a**) UMAP of cells collected from brain tissue of sham-operated mice or peri-infarct regions 7 days after stroke onset. OPCs: oligodendrocyte precursor cells, BECs: brain endothelial cells, VSMCs: Vascular smooth muscle cells. (**b**) The heatmap shows the number of differentially expressed genes (DEGs) in each cellular population when compared between the indicated mouse groups. (**c**,**d**) Comparison of total standardized read counts of recovery-associated genes in all excitatory neurons (**c**) or oligodendrocytes (**d**) on day 7 after stroke onset among the indicated mouse groups. Percentages indicate the ratio of cells showing the score of recovery-associated gene expression (dashed lines: > 0.15 for excitatory neurons (**c**), > 0.10 for oligodendrocytes (**d**)). (**e**,**f**) PCA (**e**) and heatmap (**f**) of module scores calculated by BALSAMICO analysis, which characterized the function of myeloid cells in each mouse group. (**g**) The volcano plot shows the differences in gene expression between the post-ischemic day 7 myeloid cells of PT-WT and PT-*App^NL-G-F/NL-G-F^* mice. The lower panel shows the gene ontology analysis of the genes whose expression levels significantly increased in PT-*App^NL-G-F/NL-G-F^* mice compared to PT-WT mice. (**h**) UMAPs show the differences in myeloid cell clusters of each mouse group (1454, 1334, 1727, 3441 cells for Sham-WT, Sham-*App^NL-G-F/NL-G-F^*, PT-WT, PT-*App^NL-G-F/NL-G-F^*, respectively). (**i**) Bubble heatmap chart showing the read ratio of genes that belong to gene ontologies associated with inflammatory response. The size of the circles indicates the percentage of each myeloid cell cluster that belongs to microglia (MG), infiltrating myeloid cells (IMs), and border-associated macrophage (BAM).

To compare gene expression profiles in myeloid cells, we performed a BALSAMICO analysis, which illustrates differences in cellular functions using module scores^23^. These modules are matrix variables reflecting gene expression levels in cells influenced by environments, and the enrichment of gene ontologies that belonged to each module is shown in **Supplementary** Fig.2. PCA revealed apparent differences in module scores representing cellular functions among myeloid cells of 2-month-old *App^NL-G-F^* KI and WT mice without stroke and 7 days after ischemic stroke onset (**Fig.3e**). Consistent with PCA, we found several characteristic modules (e.g., modules d, e, m, n, t, and w) unique to myeloid cells of *App^NL-G-F^* KI mice after ischemic stroke (**Fig.3f**), which were quite different from the additive combination of myeloid cell characteristics seen in the sham-operated *App^NL-G-F^* KI mice and WT mice after ischemic stroke. Thus, a unique myeloid immunity was induced in the ischemic brain with cerebral Aβ accumulation, which differed from that seen in ischemic stroke or Alzheimer’s disease solely.

Compared with that of WT myeloid cells after ischemic stroke, gene expression associated with immune response and cytokine production was increased in myeloid cells of 2-month-old *App^NL-G-F^* KI mice after ischemic stroke (**Fig.3g**). Uniform manifold approximation and projection (UMAP) of myeloid cells showed that microglial cells (MG: clusters 1−7) in sham-operated WT brain considerably changed their characteristics in sham-operated *App^NL-G-F^* KI mice (**Fig.3h**). In WT and *App^NL-G-F^* KI mice, infiltrating myeloid cells (IMs: clusters 8−10) appeared after ischemic stroke compared with sham-operated WT myeloid cell clusters (**Fig.3h**). The annotation of myeloid cell clusters including border-associated macrophage (BAM) was confirmed by marker gene expression (**Supplementary** Fig.3). When expression levels of genes included in ontologies associated with inflammatory response (gene list is shown in **Supplementary data file1**) were compared, inflammatory gene expression was observed in MG of sham-operated *App^NL-G-F^* KI mice (**Fig.3i**). Although ischemic stroke induced inflammatory responses in MG and IMs, inflammatory responses after ischemic stroke were enhanced in almost all myeloid clusters of *App^NL-G-F^* KI mice (**Fig.3i**). Thus, a unique myeloid immunity enhancing inflammatory responses was induced in the ischemic brain with Aβ accumulation.

### Anti-Aβ antibody administration restored stroke recovery in *App^NL-G-F^* KI **mice**

Next, we examined whether anti-Aβ antibody administration could prevent this pro-inflammatory myeloid immunity and restore neural repair in *App^NL-G-F^* KI mice after ischemic stroke. We administered mAducanumab to 2-month-old heterozygous *App^NL-G-F^* KI mice 24 hours after ischemic stroke onset and investigated gene expression profiles in various brain cells by scRNA-seq on day 7 after stroke onset (**Fig.4a**). Comparing expression between control-antibody- and mAducanumab-administered mice, most DEGs were detected in excitatory neurons and myeloid cells (**Fig.4b**). In excitatory neurons, recovery-associated genes whose expression was impaired in *App^NL-G-F^* KI mice (shown in **Fig.1e**) were significantly increased in mAducanumab-administered mice (**Fig.4c**,**d**). UMAP of excitatory neurons also showed increased expression of *Rgs4*, *Adcyap1*, and *Cbln4* (**Fig.4e**); these genes are important for neurotransmission, neuronal function, axon guidance, and stroke recovery^24–27^.

**Figure 4.**
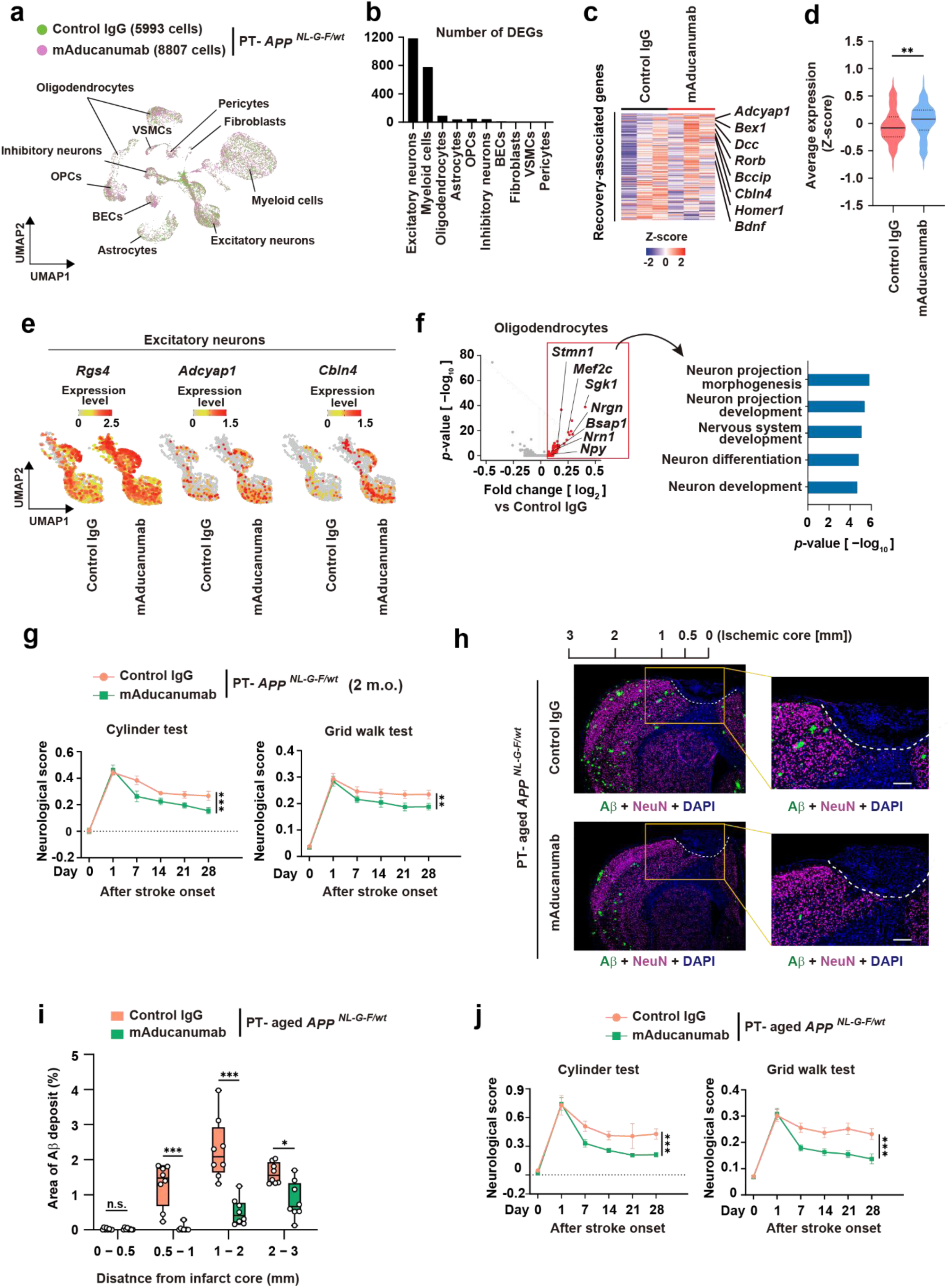
Anti-Aβ antibody resolved the impairment of stroke recovery due to cerebral Aβ accumulation. (**a**) UMAP of cells collected from peri-infarct regions 7 days after stroke onset in control-antibody- and mAducanumab-administered 2-month-old *App^NL-G-F/wt^* mice. (**b**) The number of DEGs in each cellular population compared between control-antibody- and mAducanumab-administered *App^NL-G-F/wt^* mice. (**c**,**d**) Heatmap of expression levels (**c**) and comparison of averaged expression (**d**) of recovery-associated genes in the post-ischemic day 7 peri-infarct neurons analyzed by bulk RNA-seq (n = 3 for each group). (**e**) Comparison of recovery-associated gene heatmaps in post-ischemic day 7 peri-infarct excitatory neurons between control-antibody- and mAducanumab-administered *App^NL-G-F/wt^* mice. (**f**) Volcano plot shows the differences in gene expression between the post-ischemic day 7 oligodendrocytes of control-antibody- and mAducanumab-administered *App^NL-G-F/wt^* mice. The right panel shows gene ontology analysis of the top 200 genes whose expression levels increased in mAducanumab-administered *App^NL-G-F/wt^* mice compared to control-antibody-administered *App^NL-G-F/wt^* mice. (**g**) The comparison of neurological deficits between 2-month-old (2 m.o.) *App^NL-G-F/wt^* mice administered control IgG and mAducanumab 24 hours after ischemic stroke onset (n = 8 for each group). (**h**,**i**) Immunohistochemistry of Aβ by using 82E1 in the post-ischemic day 28 brain tissue of control-antibody- or mAducanumab-administered aged *App^NL-G-F/wt^* mice (bar: 200 μm) (**h**), cerebral Aβ accumulation level in each cortical region away from the ischemic core as specified in **Fig.4h** (**i**). (**j**) The comparison of neurological deficits between the aged *App^NL-G-F/wt^* mice administered control IgG and mAducanumab 24 hours after ischemic stroke onset (n = 11 for control IgG, n = 15 for mAducanumab). \**p* < 0.05, \*\**p* < 0.01, \*\*\**p* < 0.001 vs. control-IgG-administered mice (**d**,**g**,**i**,**j**) (Mann-Whitney U test [**d**], two-way ANOVA [**g**,**j**], two-way ANOVA with Sidak’s multiple comparison test [**i**]). Error bars represent the mean ± standard error of the mean (SEM). n.s. : not significant.

In oligodendrocytes, gene ontologies associated with neuron projection and nervous system development were enriched in genes whose expression was significantly increased in mAducanumab-administered mice (**Fig.4f**). These results revealed that cerebral Aβ accumulation in *App^NL-G-F^* KI mice impaired the neural repair necessary for stroke recovery.

We then investigated neurological deficits after ischemic stroke in 2-month-old and aged heterozygous *App^NL-G-F^* KI mice. Administration of mAducanumab 24 hours after ischemic stroke onset significantly improved neurological deficits compared to the control antibody in 2-month-old heterozygous *App^NL-G-F^* KI mice (**Fig.4g**), although infarct volume was not significantly different between the two groups. When we administered mAducanumab to WT mice 24 hours after ischemic stroke onset, we did not observe an improvement in neurological deficits despite the increased Aβ level detected in post-ischemic day 7 brain tissue (**Supplementary Fig.4a**,**b**), suggesting that cerebral accumulation of aggregated Aβ before stroke onset caused the impairment of stroke recovery.

In aged heterozygous *App^NL-G-F^* KI mice, we found that mAducanumab administration efficiently removed Aβ deposits around the infarct region (**Fig.4h**,**i**). Similar to 2-month-old *App^NL-G-F^* KI mice, we did not observe any significant difference in neuronal loss around the infarct region between control-antibody- and mAducanumab-administered mice on day 28 after stroke onset (**Supplementary Fig.4c**). Neurological deficits after ischemic stroke in aged heterozygous *App^NL-G-F^* KI mice were significantly improved by mAducanumab administration 24 hours after stroke onset (**Fig.4j**). These results demonstrated the ability of anti-Aβ antibody administration within 24 hours after stroke onset to restore the neural repair necessary for stroke recovery in the brain with Aβ accumulation.

### Anti-Aβ antibody resolved pro-inflammatory myeloid immunity in the ischemic brain with Aβ accumulation

UMAPs of myeloid cells on day 7 after ischemic stroke onset did not differ between control-antibody- and mAducanumab-administered 2-month-old heterozygous *App^NL-G-F^* KI mice (**Fig.5a**). However, gene ontologies associated with inflammatory responses and IL-1β production were enriched in DEGs between the two groups (**Fig.5b**). We compared the expression levels of genes included in ontologies associated with inflammatory response in each myeloid cluster and found that inflammatory responses were resolved in almost all the post-ischemic day 7 myeloid clusters by mAducanumab administration 24 hours after stroke onset (**Fig.5c**). Thus, mAducanumab resolved the pro-inflammatory myeloid immunity to restore neuronal repair after ischemic stroke.

**Figure 5.**
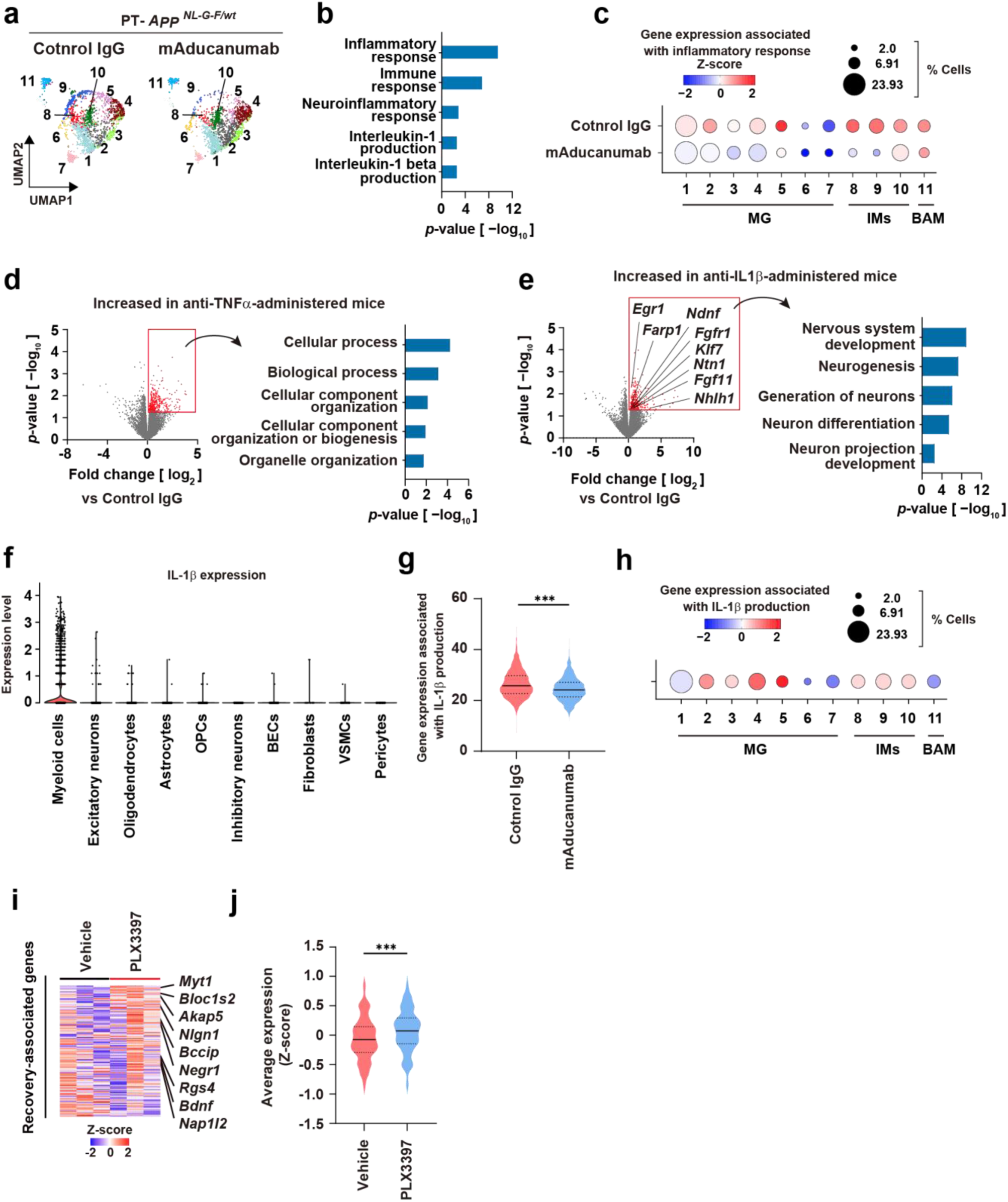
Post-stroke anti-Aβ antibody administration prevented the pro-inflammatory myeloid immunity impeding neuronal repair. (**a**) UMAPs show the myeloid cell clusters of control-antibody- or mAducanumab-administered 2 m.o. *App^NL-G-F/wt^* mice 7 days after ischemic stroke onset. (**b**) Results of gene ontology analysis of the DEGs whose expression levels significantly decreased in mAducanumab-administered *App^NL-G-F/wt^* mice. (**c**) Bubble heatmap chart showing the read ratio of genes that belong to gene ontologies associated with the inflammatory response. The size of the circles indicates the percentage of each myeloid cell cluster. (**d**,**e**) Volcano plots show the differences in gene expression between post-ischemic day 7 peri-infarct neurons of *App^NL-G-F/NL-G-F^* mice administered control IgG and anti-TNFα antibody (**d**) or anti-IL1β antibody (**e**) 24 hours after stroke onset (n = 3 for each group). Each right panel shows the gene ontology analysis of the genes whose expression levels significantly increased in mice administered the antibody against cytokine. (**f**) mRNA expression levels of *Il1b* in each cell collected from post-ischemic day 7 brain of *App^NL-G-F/wt^* mice administered control IgG. (**g**,**h**) Comparison of gene expression included in gene ontology “Interleukin-1 beta production” in the post-ischemic day 7 myeloid cells of *App^NL-G-F/wt^* mice (2056 cells for control IgG, 1936 cells for mAducanumab) (**g**) and in each myeloid cluster of *App^NL-G-F/wt^* mice administered control IgG (**h**). (**i**,**j**) Bulk RNA-seq results of post-ischemic day 7 peri-infarct neurons in *App^NL-G-F/NL-G-F^* mice administered vehicle or PLX3397 (n = 3 for each group) for 21 consecutive days before the induction of ischemic stroke. Heatmap shows the expression levels of recovery-associated genes (**i**). Averaged expression levels of each recovery-associated gene are compared by violin plot (**j**). \*\*\**p* < 0.001 vs. control-IgG-administered mice (**g**) or vehicle-administered mice (**j**) (Mann-Whitney U test [**g**,**j**]).

To clarify the important inflammatory mediators that impaired neural repair in the ischemic brain with Aβ accumulation, we examined whether the administration of antibodies against inflammatory cytokines restored gene expression associated with recovery processes in the peri-infarct neurons by bulk RNA-seq. The administration of anti-TNFα antibody to 2-month-old homozygous *App^NL-G-F^* KI mice 24 hours after stroke onset did not alter gene expression associated with stroke recovery in peri-infarct neurons on day 7 after stroke onset (**Fig.5d**). The administration of anti-IL1β antibody significantly increased gene expression associated with neural repair in peri-infarct neurons (**Fig.5e**), revealing that enhanced production of IL-1β was a pivotal factor in causing stroke recovery impairment by cerebral Aβ accumulation.

A major source of IL-1β was myeloid cells in the ischemic brain with Aβ accumulation (**Fig.5f**). The expression levels of genes in myeloid cells included in the “IL-1β production” ontology were significantly decreased by mAducanumab administration 24 hours after stroke onset (**Fig.5g**). Gene expression included in IL-1β production was predominantly detected in MG clusters rather than IMs and BAM (**Fig.5h**). To clarify whether the existence of pro-inflammatory microglial cells before stroke onset in *App^NL-G-F^* KI mice (as shown in **Fig.3i**) was pivotal in stroke recovery impairment, we examined administration of PLX3397, an inhibitor of colony stimulating factor 1 receptor (CSF1R), to deplete microglia from brain before stroke onset^28^. We found that microglial depletion before the induction of ischemic stroke in homozygous *App^NL-G-F^* KI mice significantly increased recovery-associated gene expression in peri-infarct neurons on day 7 after stroke onset (**Fig.5i**,**j**), revealing that cerebral accumulation of aggregated Aβ induced pro-inflammatory microglia before stroke onset to impair stroke recovery.

### TLR2 and TLR4 were major signaling pathways to impair stroke recovery in ***App^NL-G-F^* KI mice**

In Alzheimer’s disease or ischemic stroke, Aβ and damage-associated molecular patterns (DAMPs) activate Toll-like receptor 2 (TLR2) and TLR4 in myeloid and brain cells to trigger intracerebral sterile inflammation^29–32^. We investigated whether TLR2 and TLR4 were major signaling pathways to induce the unique myeloid immunity that impaired stroke recovery in the ischemic brain with Aβ accumulation. *App^NL-G-F^* KI mice were crossed with *Tlr2*^−/−^; *Tlr4*^−/−^ (TLR2/4 DKO) mice, and gene expression profiles in various cells on day 7 after ischemic stroke onset were analyzed using scRNA-seq (**Fig.6a**). DEGs between *App^NL-G-F^* KI mice and TLR2/4-double-deficient *App^NL-G-F^* KI mice were strongly detected in myeloid cells, oligodendrocytes, and excitatory neurons (**Fig.6b**). In myeloid cells, gene ontologies associated with inflammatory responses were enriched in DEGs (**Fig.6c**). Although UMAP of myeloid cells was little altered between the two groups (**Fig.6d**), gene expression included in the ontologies associated with inflammatory response was apparently decreased in almost all myeloid clusters (**Fig.6e**). This confirmed that the TLR2 and TLR4 signaling pathways in brain cells and infiltrating immune cells were essential for pro-inflammatory myeloid immunity formation in the ischemic brain with Aβ accumulation. Indeed, PCA showed that gene expression profiles of post-ischemic day 7 myeloid cells in TLR2/4-double-deficient *App^NL-G-F^* KI mice were closer to those of post-stroke WT mice than to those of post-stroke *App^NL-G-F^* KI mice (**Fig.6f**).

**Figure 6.**
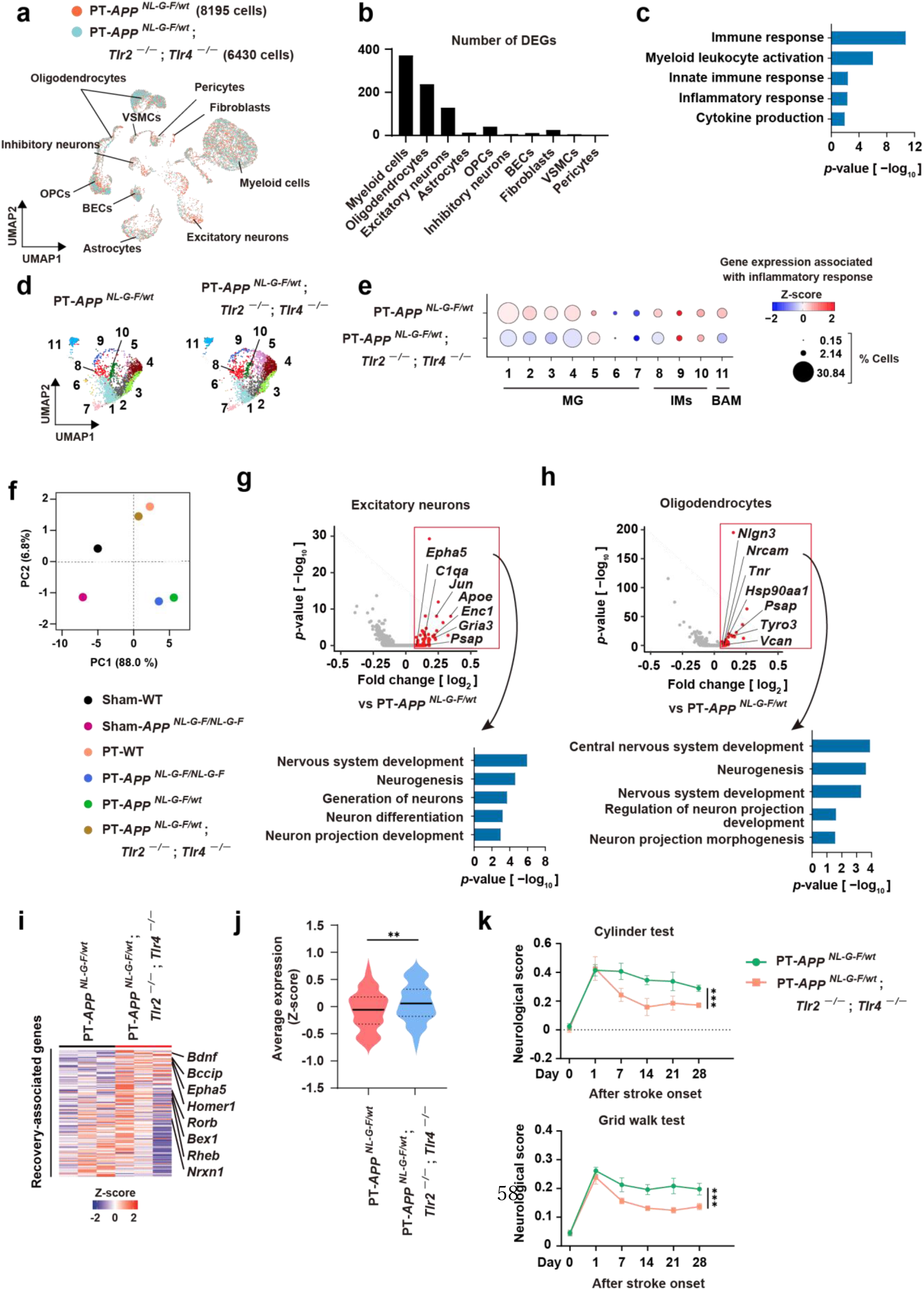
Cerebral Aβ accumulation impaired stroke recovery through TLR2 and TLR4 activation. (**a**) UMAP of cells collected from peri-infarct regions 7 days after stroke onset in 2-month-old *App^NL-G-F/wt^* mice and *Tlr2*^−/−^; *Tlr4*^−/−^; *App^NL-G-F/wt^* mice. (**b**) The number of DEGs in each cellular population between *App^NL-G-F/wt^* mice and *Tlr2*^−/−^; *Tlr4*^−/−^; *App^NL-G-F/wt^* mice. (**c**) Results of gene ontology analysis of the DEGs whose expression levels decreased significantly in myeloid cells of *Tlr2*^−/−^; *Tlr4*^−/−^; *App^NL-G-F/wt^* mice compared to *App^NL-G-F/wt^* mice. (**d**) UMAPs show the myeloid cell clusters of *App^NL-G-F/wt^* mice or *Tlr2*^−/−^; *Tlr4*^−/−^; *App^NL-G-F/wt^* mice. (i) The read ratio of genes belonging to gene ontologies associated with the inflammatory response in each myeloid cluster. Circle size indicates the percentage of each myeloid cell cluster. (**f**) PCA of gene expression profiles that belong to the gene ontologies associated with inflammatory response in the myeloid cells of each mouse group. Myeloid cells of PT-mice were isolated 7 days after stroke onset. (**g**,**h**) Volcano plots show differences in gene expression between post-ischemic day 7 excitatory neurons (**g**) or oligodendrocytes (**h**) of *App^NL-G-F/wt^* mice and *Tlr2*^−/−^; *Tlr4*^−/−^; *App^NL-G-F/wt^* mice. The lower panel shows the gene ontology analysis of the top 200 genes whose expression levels increased in *Tlr2*^−/−^; *Tlr4*^−/−^; *App^NL-G-F/wt^* mice compared to *App^NL-G-F/wt^* mice. (**I**,**j**) Bulk RNA-seq result of post-ischemic day 7 peri-infarct neurons in 2-month-old *App^NL-G-F/wt^* mice and *Tlr2*^−/−^; *Tlr4*^−/−^; *App^NL-G-F/wt^* mice (n = 3 for each group). The heatmap shows the expression levels of recovery-associated genes Comparison of averaged expression levels of each recovery-associated gene (j) (**k**) Time-dependent changes of neurological deficits between 2-month-old *App^NL-G-F/wt^* mice and *Tlr2*^−/−^; *Tlr4*^−/−^; *App^NL-G-F/wt^* mice (n = 8 for *App^NL-G-F/wt^* mice, n = 5 for *Tlr2*^−/−^; *Tlr4*^−/−^; *App^NL-G-F/wt^* mice). \*\**p* < 0.01, \*\*\**p* < 0.001 vs. PT-*App^NL-G-F/wt^* mice (**j**,**k**) (Mann-Whitney U test [**j**], two-way ANOVA [**k**]). Error bars represent the mean ± standard error of the mean (SEM).

TLR2/4-double deficiency in *App^NL-G-F^* KI mice significantly increased gene expression associated with stroke recovery processes, such as nervous system development and neuron projection development, in excitatory neurons and oligodendrocytes around infarct regions on day 7 after stroke onset (**Fig.6g**,**h**). When we evaluated gene expression profiles in neurons isolated from the peri-infarct region by bulk RNA-seq, the expression of recovery-associated genes (shown in **Fig.1e**) was significantly increased by TLR2/4-double deficiency in *App^NL-G-F^* KI mice on day 7 after stroke onset (**Fig.6i**,**j**). TLR2/4-double deficiency significantly improved neurological deficits after ischemic stroke in *App^NL-G-F^* KI mice (**Fig.6k**). Thus, TLR2 and TLR4 signaling pathways were critical in impaired stroke recovery in the ischemic brain with Aβ accumulation.

## Discussion

In this study, cerebral Aβ accumulation induced pro-inflammatory microglia with little changes in neuronal gene expression before ischemic stroke onset. No deficits in synaptic plasticity were reported in 2-month-old *App^NL-G-F^* KI mice^33^. It is possible that similar phenomena occur in the Aβ-accumulated brains of patients with preclinical Alzheimer’s disease whose frequency is 10−20% at age 50−70 years, 30−65% at age over 70 years^34–37^. A week after ischemic stroke onset, inflammation by myeloid cells was almost resolved, and remarkable recovery mechanisms were triggered in peri-infarct neurons; however, these mechanisms were considerably dysfunctioned even by slight cerebral Aβ accumulation, leading to poor functional prognosis after stroke. Through TLR2 and TLR4 activation in brain cells and infiltrating immune cells, a unique pro-inflammatory myeloid immunity— distinct from that seen in either Alzheimer’s disease or ischemic stroke alone—was induced by cerebral Aβ accumulation in the ischemic brain, resulting in impaired stroke recovery (**Supplementary** Fig.5). Mathematical modeling analysis confirmed that an original factor resulting from ischemic stroke comorbid with Alzheimer’s disease better explained the formation of this unique myeloid immunity than simply additive effects of both. Although disease-specific glial and myeloid cells have recently attracted attention^38,39^, we thus discovered the unique myeloid immunity formation in combined neurological disorders that impeded neurological recovery.

When cerebrovascular disease and Alzheimer’s disease coexist, each progressively worsens the pathology of the other, resulting in a vicious cycle. This leads to poor functional prognosis, making it difficult to develop therapeutic drugs for stroke and neurocognitive disorders^40^. A long-time research goal has been to clarify whether stroke recovery mechanisms and long-term prognosis are affected by spontaneous cerebral accumulation of pathogenic Aβ, because evaluating long-term stroke recovery by exogenous Aβ peptide injection or APP protein overexpression using *App* Tg mice has been difficult. In fact, some reports, but not others, showed exacerbated neuronal loss or neurological deficits by Aβ injection^41–50^ or APP overexpression^17,51–54^ due to unpredicted artificial effects rather than spontaneous cerebral Aβ accumulation. Although one report shows that Aβ peptide injection increases the cells producing insulin-like growth factor 1 (IGF1)^55^, a typical neurotrophic factor, we did not observe any alteration of reparative functions in myeloid cells in the current study.

Cerebral Aβ accumulation after ischemic stroke is well known, and this worsens Alzheimer’s disease pathologies^56–60^. Stroke-induced cerebral Aβ accumulation may disrupt the vascular function of maintaining cerebral blood flow. Although one report considered whether preventing stroke-induced cerebral Aβ accumulation is beneficial for avoiding poor functional prognosis^61^, we could not demonstrate a long-term benefit of anti-Aβ antibody administration to a murine model of ischemic stroke without cerebral Aβ accumulation. This can be explained by a previous report demonstrating promoted myelin repair by stroke-induced cerebral Aβ accumulation^42^.

Administration of anti-Aβ antibodies causes amyloid-related imaging abnormalities with edema or effusion^10^; however, we did not observe any signs of exacerbated cerebral inflammation, intracerebral hemorrhage, or neurotoxicity when anti-Aβ antibodies were administered after ischemic stroke onset. Anti-Aβ antibody resolved the myeloid inflammation impeding neural repair and restored stroke recovery in ischemic brain with slight or remarkable Aβ accumulation. Suppression of IL-1β was pivotal for restoring neural repair, which was consistent with a previous *in vitro* study^62^. Thus, post-stroke aducanumab administration was beneficial for ischemic stroke with cerebral Aβ accumulation to improve functional prognosis. We predict that treatment using anti-Aβ antibodies to remove cerebral Aβ in patients with Alzheimer’s disease may be useful for preventing poor functional prognosis due to ischemic stroke onset.

## Acknowledgments

We thank Mrs. Yoshiko Yogiashi and Kumiko Kurabayashi for their experimental assistance throughout the study. We appreciate Mrs. Mariko Yoshimura for her assistance with preparing the manuscript, and Dr. Erika Seki for her kind cooperation in preparing human brain sections. This work was supported by CREST from AMED under grant number JP23gm1210010 (T.S.); a Grant-in-Aid for Scientific Research on Innovative Areas (Dynamic Regulation of Brain Function by the Glia decoding [23H04181]) from the Ministry of Education, Culture, Sports, Science and Technology of Japan (MEXT) (T.S.); JSPS KAKENHI Grants-in-Aid for Scientific Research (A) (24H00626) (T.S.); a Toray Science and Technology Grant (T.S.); the Takeda Science Foundation (T.S., S.S., J.T.); the Uehara Memorial Foundation (S.S.); Research Foundation for Opto-Science and Technology (S.S.); and grants from the MSD Life Science Foundation (T.S.), Senri Life Science Foundation (T.S.), Ono Pharmaceutical Foundation for Oncology, Immunology, and Neurology (T.S.); Charitable Trust Mihara Cerebrovascular Disorder Research Promotion Fund (T.S.); Medical Research Center Initiative for High Depth Omics, TMDU, Nanken-Kyoten, TMDU, Multilayered Stress Diseases (JPMXP1323015483), TMDU.

## Author contributions

K.O. designed and performed the experiments, analyzed the data, and wrote the manuscript; J.T., S.S. and K.H. provided technical advice and analyzed the data; Y.H. and H.K. analyzed the RNA-seq data and participated in discussion; K.A. and T.S. analyzed the scRNA-seq results and performed module analysis; I.K. and T.K. provided the samples and advice in the analysis of human brain; T.S. kindly provided experimental advice to the entire study using *App* KI mice; T.S. initiated and directed the entire study, designed the experiments, and wrote the manuscript.

## Materials and Methods

### Mice

All mice were maintained in a conventional facility at the Tokyo Medical and Dental University or Tokyo Metropolitan Institute of Medical Science in Tokyo, Japan. App^NL-G-F/NL-G-F^ mice^20^ (C57BL/6J background) were kindly provided by Dr. Takashi Saito (Nagoya City University) and Dr. Takaomi Saido (RIKEN Brain Science Institute). Tlr2^−/−^; Tlr4^−/−^ mice^63^ (C57BL/6J background) were kindly provided by Dr. Shizuo Akira (Osaka University) to be crossed with App^NL-G-F/NL-G-F^ mice. C57BL/6J mice were used for WT mice. All experiments were approved by the Institutional Animal Research Committee of the Tokyo Medical and Dental University (approval number: A2023-172C2) and Tokyo Metropolitan Institute of Medical Science (approval number: 21-004).

### Mouse model of ischemic stroke

Male mice were randomly selected for experiments. Mice aged over 16 months were used for aged mouse experiments. Five to fifteen mice were estimated to be necessary to reach sufficient statistical power. To generate the photothrombotic ischemia model, mice were anesthetized with isoflurane in a mixture of 70% nitrous oxide and 30% oxygen. Body temperature was maintained at 35°C during surgery. Rose Bengal (Sigma-Aldrich) was intravenously injected at a dose of 20 mg/kg body weight and activated by focal illumination at the primary motor cortex (1.5 mm lateral and 0.2 mm anterior to bregma in the left hemisphere) with a green laser (561 nm, 4 mW, Coherent Inc.) for 5 minutes. After the scalp was sutured, the mice were allowed to recover from anesthesia. Although no mice were dead after surgery or during the phase for the assessments of neurological deficits, the comparison of survival rate in each experiment were shown in **Supplementary** Fig.6.

### Assessments of neuronal injury after ischemic stroke

To evaluate long-term neurological function, we performed the cylinder test and the grid walk test on day 0 (before induction of ischemic stroke) and on days 1, 7, 14, 21, and 28 after ischemic stroke onset. For the cylinder test, mice were placed in a clear cylinder and the number of times the forelimbs contacted the wall was recorded for 5 minutes. Neurological score was calculated as (number of nonimpaired forelimb contacts − number of impaired forelimb contacts)/(number of nonimpaired forelimb contacts + number of impaired forelimb contacts). For the grid walk test, mice were placed on a grid surface and allowed to move across it. Before induction of ischemic stroke, mice were trained to walk freely for 10 minutes per day on the grid surface for 2 days to familiarize them with it. The number of foot-slips for the impaired forelimb and the total footsteps were recorded for 3 minutes. Neurological score was calculated as (number of foot-slips for impaired forelimb)/(number of total footsteps) To evaluate infarct volume, mice were euthanized with deep sedation and then transcardially perfused with phosphate-buffered saline (PBS) followed by 4% paraformaldehyde/PBS. Forebrains were removed and fixed with 4% paraformaldehyde overnight at 4°C. Fixed forebrains coronally sliced at 1 mm thickness were embedded in paraffin. The infarct area was detected by the immunohistochemistry of microtubule-associated protein 2 (MAP2) and measured using ImageJ software (National Institutes of Health, Bethesda, MD, USA).

### Treatments with mAducanumab

Hybridoma producing control IgG antibody and the plasmids encoding mouse chimeric aducanumab (mAducanumab) against human Aβ oligomer or fibrils were kindly provided by Prof. David M. Holtzman (Washington University). The methods for generating mAducanumab were previously described elsewhere^64^. Briefly, plasmids encoding mAducanumab were co-transfected in FreeStyle™ 293-F cells (ThermoScientific). mAducanumab was purified from cultured medium by using protein G sepharose beads (Cytiva). Two hundred micrograms of control IgG or mAducanumab was intravenously administered to mice 24 hours after ischemic stroke onset.

### Whole-cell patch clamp recording

WT or App^NL-G-F/NL-G-F^ mice 4 weeks after ischemic stroke onset were used for patch clamp experiments. The mice were deeply anesthetized with isoflurane and transcardially perfused with ice-cold N-methyl-D-glutamine (NMDG)-artificial cerebrospinal fluid (NMDG-ACSF; 93 mM NMDG, 20 mM HEPES, 2.5 mM KCl, 30 mM NaHCO_3_, 1.2 mM NaH_2_PO_4_, 0.5 mM CaCl_2_, 10 mM MgCl_2_, 25 mM glucose, 1 mM kynurenic acid, 0.2 mM lidocaine HCl, 5 mM sodium ascorbate, 3 mM sodium pyruvate, 2 mM thiourea, bubbled with 95% O_2_ and 5% CO_2_, pH 7.4). The brain was removed and coronally sliced at 300 μm thickness in NMDG-ACSF using a micro slicer (DTK-1000N, Dosaka EM). The brain slices 1.0-1.5 mm anterior to the ischemic core were collected and incubated in NMDG-ACSF at 37°C for 15 min, followed by incubation in HEPES-ACSF (92 mM NaCl, 20 mM HEPES, 2.5 mM KCl, 30 mM NaHCO_3_, 1.2 mM NaH_2_PO_4_, 0.5 mM CaCl_2_,10mM MgCl_2_, 25 mM glucose, 5 mM sodium ascorbate, 3 mM sodium pyruvate, 2 mM thiourea, bubbled with 95% O_2_ and 5% CO_2_, pH 7.4) at room temperature for longer than 1 hour before recording.

After incubation, the brain slices were placed in a recording chamber perfused with ACSF (125 mM NaCl, 2.5 mM KCl, 25 mM NaHCO_3_, 1.25 mM NaH_2_PO_4_, 2 mM CaCl_2_, 1 mM MgSO_4_, 11 mM glucose, bubbled with 95% O_2_ and 5% CO_2_, pH 7.4) at 30°C. For recordings of miniature excitatory post-synaptic currents (mEPSCs), 1 μM tetrodotoxin was added to ACSF. Neurons were visualized using an infrared differential interference contrast (IR-DIC) microscope (FN1, Nikon). Patch pipettes (3-5 MΩ) were made of glass capillaries (GC150F, Harvard Apparatus) and filled with an internal solution (130 mM Cs-gluconate, 8 mM CsCl, 0.2 mM EGTA, 10 mM HEPES, 1 mM MgCl_2_, 3 mM MgATP, 0.3 mM Na_2_GTP, 10 mM Na_2_-phosphocreatine, 0.3% Biocytin, pH 7.2). Whole-cell patch clamp recordings were obtained from layer 2/3 pyramidal cells in the motor cortex anterior to the infarct area. mEPSCs were recorded in voltage-clamp configuration at a holding potential of −70 mV for 10 min. Current signals were amplified and digitized at 20 kHz using a patch clamp amplifier (EPC10, HEKA) and stored using Patchmaster software (HEKA). Data were filtered at 1 kHz by low-pass Butterworth filter. Current changes larger than 5 pA were considered as mEPSC. Data were discarded when series resistance exceeded 25 MΩ.

### Isolation of brain cells for next-generation sequencing

Brain cells including neurons were isolated from adult ischemic brain tissue according to the experimental procedure previously reported elsewhere^15^. Mice under deep anesthesia were perfused transcardially with a sufficient volume of NMDG-ACSF. The forebrains were removed and soaked in NMDG-ACSF on ice. The removed forebrains were sliced into 300 μm thick sections. The brain sections were incubated in NMDG-ACSF for 15 min at 34°C. The peri-infarct brain tissues that were considered to be expanded 0.5−2 mm lateral from the ischemic core were cut out from the brain slices. The collected peri-infarct brain tissues were digested with pronase in HEPES-ACSF bubbled with 95% O_2_/5% CO_2_ for 70 min. Digested peri-infarct brain tissues were triturated with fire-polished Pasteur pipettes after three washes with HEPES-ACSF containing 1% fetal bovine serum (FBS) and 0.2% bovine serum albumin (BSA). The dissociated brain cells were stained with anti-Thy1.2 PE (53-2.1, BioLegend), anti-ACSA2 APC (130-116-142, Miltenyi Biotec), anti-CD31 APC (390, BioLegend) or anti-CD31 PE-Cy7 (390, BioLegend), anti-CD45 APC (30-F11, BioLegend) or anti-CD45 FITC (30-F11, eBioscience), anti-O4 APC (130-118-978, Miltenyi Biotec), anti-CD140a APC (APA5, BioLegend), and anti-CD140b APC (APB5, BioLegend), and analyzed by FACS. The Thy1.2-positive and ACSA2-, CD31-, CD45-, O4-, CD140a-, and CD140b-negative population was confirmed as neuronal cells, as previously described^15^, and isolated for RNA-seq analysis. SYTOX Green nucleic acid stain (S7020, Invitrogen) was added to remove dead brain cells for scRNA-seq analysis.

### RNA-seq

Total RNA was purified using RNeasy Mini Kit (Qiagen) after the neuronal cell-enriched population isolated from normal or peri-infarct brain tissue was pelleted for 5 min at 300×g at 4°C. To prepare RNA-seq libraries from total RNA, Ovation SoLo RNA-Seq Library Preparation Kit (Tecan) was used according to the manufacturer’s protocol. The libraries were sequenced by Illumina HiSeq after the amount and fragment size distribution of obtained cDNA libraries were examined using an Agilent 2100 Bioanalyzer. After conducting Trimmomatic to remove adapter and low-quality sequences, reads were aligned to mm10 (annotation file) by hisat2, and the read counts for each gene were determined by featureCounts. Differentially expressed genes were identified by edgeR (glmQLFTest function). The integrated data of replicated samples from three independent experiments after the trimmed mean of M-values (TMM) normalization were used to draw Volcano plots. Gene ontology analysis was performed on genes whose expression levels were significantly increased or decreased in the comparison between two groups by g:Profiler (version e109_eg56_p17_773ec798) with g:SCS multiple testing correction methods applying a significance threshold of 0.05. To set the background genes, all genes expressed in neurons (TPM > 0) were used. Recovery-associated genes defined as genes comprising the gene ontology terms “Nervous system reconstruction”, “Synaptic organization”, and “Remyelination” (as previously described^15^) whose expression was significantly increased in the neurons of WT mice after ischemic stroke.

### scRNA-seq

scRNA-seq libraries were prepared using Chromium Next GEM Single-cell 3’ Reagent Kits v3.1 (10x Genomics) according to the manufacturer’s protocol. Single cells were loaded onto chromium chips with a capture target of 2000 neurons and 8000 non-neuronal brain cells per sample. Libraries were sequenced on a DNBSEQ-G400 (MGI) with a targeted sequencing depth of more than 50,000 read pairs per cell. Read mapping and counting were processed with the 10x Genomics Cell Ranger 6.0.1^65^ with the mm10 reference genome assembly and gene annotation provided by 10x Genomics. Doublets were removed using scDbletFinder v3.0 with default parameters. The SCTransformation method incorporated in Seurat 4.3.0 was conducted to normalize UMI counts for the individual genes^66,67^. The single-cell gene expression profiles of the 10 samples (normal brain tissue of sham-operated WT or App^NL-G-F/NL-G-F^ mice, two replicates of day 7 post-ischemic brain tissues of WT mice and App^NL-G-F/NL-G-F^ mice, and a single day 7 post-ischemic brain tissue of App^NL-G-F/wt^ mice, Tlr2^−/−^; Tlr4^−/−^; App^NL-G-F/wt^ mice, App^NL-G-F/wt^ mice administered control IgG or mAducnaumab) were merged by the Seurat anchor-based integration method^68^. This integrated count matrix was used to cluster the cells with the shared nearest-neighbor approach implemented in Seurat. Cell populations were identified by distinguishing them by the expression of marker genes specific to each brain cell type, and cell populations with marker genes overlap were excluded manually. The UMAP plots were generated with the Seurat Dimplot module by using the normalized UMI counts. DEGs were visualized in a volcano plot. Gene ontology analysis in myeloid cells, excitatory neurons, and oligodendrocytes was performed by g:Profiler (version e109_eg56_p17_773ec798 or version e111_eg58_p18_30541362) with g:SCS multiple testing correction methods applying a significance threshold of 0.05. All genes expressed (UMI counts > 0) in each cell type were used as the background. For the analysis of gene expression associated with recovery processes after brain injury, the expression level of recovery-associated genes as previously described^15^ was scored by AddModuleScore in Seurat and visualized with ViolinPlot. For the analysis of gene expression associated with the inflammatory response (shown in **Supplementary data file 1**) or included in gene ontology “IL-1β production” in myeloid cells, the UMI counts of genes belonging to the ontologies were averaged in each myeloid cluster and then shown in a bubble plot of Z-score for comparison among all of the obtained myeloid clusters.

### Immunohistochemistry

To detect Aβ in murine brain tissue, forebrains collected from sham-operated mice or photothrombotic mice administered control IgG or mAducanumab were coronally sliced at 1-mm thickness and then embedded in paraffin. The thin-sliced sections were deparaffinized and then boiled in citrate buffer using a microwave oven. Endogenous peroxidase activity was blocked using 1% H_2_O_2_/methanol. The thin-sliced sections were then blocked for 30 minutes with a blocking buffer (TNT buffer; TNB) prepared by dissolving Blocking Reagent Powder (AKOYA Biosciences) in TNT solution (0.1% Tween 20, 0.1 M Tris/HCl (pH 8.0), 0.18 M NaCl, adjusted to pH 7.5). After blocking with TNB, the sections were incubated with anti-human Aβ (82E1, IBL) antibody or mAducanumab, anti-NeuN (ABN78, Millipore) antibody overnight at 4°C. Secondary antibodies, anti-rabbit IgG antibody conjugated with Alexa Fluor 546 (A11701, Invitrogen) and EnVision+ system-HRP Labelled Polymer Anti-mouse (DAKO) were used for detection of the primary antibody. Tyramide signal amplification was performed using TSA fluorescein Reagent (AKOYA Biosciences) according to the manufacturer’s protocol. Images of the sections were captured using a fluorescence microscope (BZ-X710, Keyence). Immunoreactive areas or the number of NeuN+ neurons at each distance from the ischemic core were quantified by ImageJ software (National Institutes of Health).

To detect Aβ in the human brain, after blocking with 1% H_2_O_2_/methanol and TNB, thin-sliced sections were incubated with anti-human Aβ (82E1, IBL) antibody. After several washes and incubation with EnVision+ system-HRP Labelled Polymer Anti-mouse (DAKO), 3,30-diaminobenzidine (DAB) Substrate Kit (Nichirei) was used for the detection of Aβ before slides were counterstained with hematoxylin (SCY). Clinical information of ischemic stroke patients is shown in **Supplementary Table 2**. Three different 0.25-mm^2^ areas were evaluated using ImageJ software (National Institutes of Health) for cerebral Aβ accumulation in the peri-infarct region and hippocampus. Ischemic stroke patients were separated into three groups as negligible (< 0.2 %), slight (< 2 %), and remarkable Aβ deposition (> 2 % of 1.5 mm^2^ area).

For the detection of BDNF^+^ or NELL1^+^ neurons of the human brain sample, the thin-sliced sections were deparaffinized and boiled in citrate buffer using a microwave oven. Sections were pretreated with PBS containing 0.1% Triton-X100 for 15 min followed by incubation with TrueBlack (Biotium) to suppress autofluorescence. After blocking with Blocking One Histo (Nacalai Tesque), the sections were incubated with anti-BDNF (ab203573, Abcam), NELL1 (ab197315, Abcam), or NeuN (MAB377, Millipore) antibody overnight at 4°C. Secondary antibodies, anti-mouse IgG antibody conjugated with Alexa Fluor 546 (A11018, Invitrogen) and anti-rabbit IgG antibody conjugated with Alexa Fluor 488 (A11070, Invitrogen), were used for detection of the primary antibody. Images of the sections were captured using a fluorescence microscope (BZ-X710, Keyence) after the treatment with TrueVIEW Autofluorescence Quenching Kit (Vector Laboratories). The number of BDNF- or NELL1-positive NeuN^+^ neurons within four 0.0625-mm^2^ areas were counted in peri-infarct region.

### Enzyme-linked immunosorbent assay (ELISA)

Insoluble materials from the mouse cortical hemisphere were dissolved in guanidine-HCl (GuHCl fraction) as previously described^69^. Aβ_x-42_ in the GuHCl fraction was quantified using an Aβ ELISA kit (Wako) according to the manufacturer’s protocol. To qualify Aβ_x-42_ carrying the Arctic mutation, synthetic human Arctic Aβ peptides (Peptide Institute) were used for generating standard curves.

### BALSAMICO analysis

BALSAMICO (Bayesian Latent Semantic Analysis of Microbial COmmunities) analysis was initially developed to evaluate the associations between microbiota and their environmental factors^23^. This analyzing strategy is also useful for detecting key factors in the associations between cells in organs and their environment.

In this study, we applied BALSAMICO to precisely analyze gene expression patterns of myeloid cells under different experimental conditions. Specifically, we set up four experimental conditions:

1. Normal brain tissue from sham-operated WT mice
2. Normal brain tissue from sham-operated *App^NL-G-F/NL-G-F^* mice
3. Brain tissue from WT mice 7 days post-ischemia
4. Brain tissue from *App^NL-G-F/NL-G-F^* mice 7 days post-ischemia

Using gene-specific UMI count data from myeloid cells collected under each condition as input, we employed BALSAMICO to extract latent gene expression modules. This method allowed us to represent the gene expression profile of individual cells as a combination of multiple modules, quantitatively evaluating the contribution of each cell and gene to the modules, as well as the impact of experimental conditions on the modules.

In this analysis, we determined the number of modules using 10-fold cross-validation. Representative genes for each module were selected based on the harmonic mean of their frequency and exclusivity^70^. To elucidate the functional significance of the selected gene sets, we performed gene enrichment analysis using Fisher’s exact test, with the IMPaLA database (http://impala.molgen.mpg.de/wsdoc) as the reference gene set^71^.

### Depletion of microglial cells by PLX3397

PLX-3397 (Pexidartinib, MedKoo Biosciences) was administered to *App^NL-G-F/NL-G-F^* mice by oral gavage for 21 consecutive days at a dose of 40 mg kg^−1^. Brain ischemia was induced by photothrombosis on the day following the final administration on day 21.

### Statistical analysis

Data are displayed as minimum to maximum box-and-whisker plots or means ± standard error of the mean (SEM). To analyze the differences between two groups, unpaired Student’s *t*-test or Mann–Whitney U test were used. To analyze the differences among three or more groups, statistical significance was determined by one-way analysis of variance (ANOVA) followed by *post hoc* multiple comparison tests (Dunnett’s test) or two-way ANOVA followed by *post hoc* multiple comparison tests (Tukey’s test or Sidak’s test). *p* < 0.05 was considered to represent a statistically significant difference.

