## Supplementary material for "Anti-amyloid beta therapy resolves stroke recovery impairment caused by Alzheimer’s disease": Supplematal data

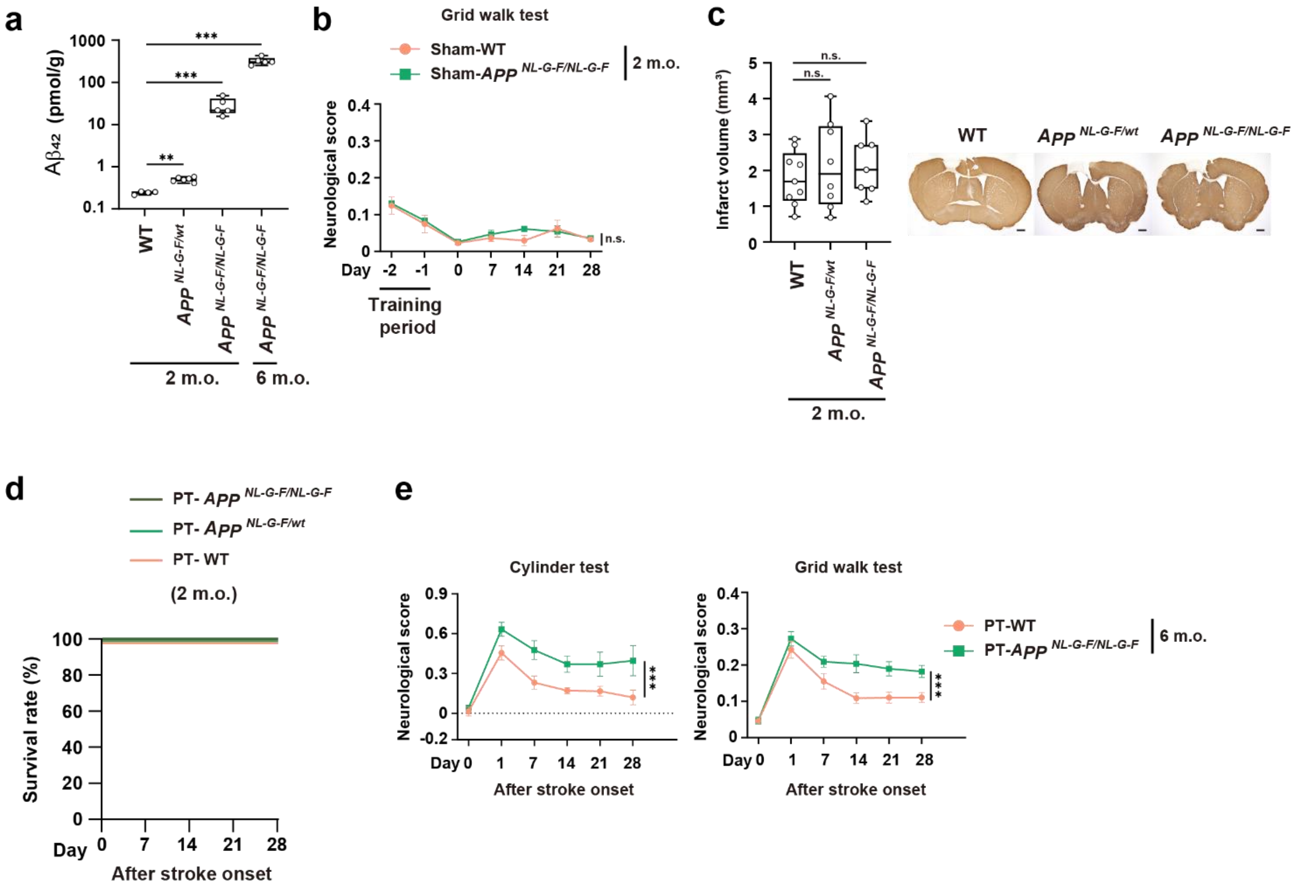


**Supplementary Figure 1.**

(**a**) Quantification of human Aβ42 peptide [pmol] in the brain tissue [g] by ELISA before the induction of ischemic stroke. (**b**) The comparison of neurological function during 2 days of the training period and subsequent 28 days of the evaluation period between two-month-old (2 m.o.) sham-operated WT mice and $App^{NL\text{-}G\text{-}F/NL\text{-}G\text{-}F}$ mice without ischemic stroke (n = 3 for each group). (**c**) The comparison of infarct volume among two-month-old WT mice, $App^{NL\text{-}G\text{-}F/wt}$ mice, and $App^{NL\text{-}G\text{-}F/NL\text{-}G\text{-}F}$ mice 28 days after ischemic stroke onset (n = 9 for WT, n = 8 for $App^{NL\text{-}G\text{-}F/wt}$, n = 7 for $App^{NL\text{-}G\text{-}F/NL\text{-}G\text{-}F}$, bar: 500 μm). (**d**) The survival rate of two-month-old WT, $App^{NL\text{-}G\text{-}F/wt}$ , and $App^{NL\text{-}G\text{-}F/NL\text{-}G\text{-}F}$ after ischemic stroke onset. (**e**) The time-dependent changes of neurological deficits after ischemic stroke onset in six-month-old (6 m.o.) mice (n = 5 for PT-WT, n = 8 for PT-$App^{NL\text{-}G\text{-}F/NL\text{-}G\text{-}F}$). $^{**}p < 0.01$, $^{***}p < 0.001$ vs. 2 m.o. WT (**a**) or PT-WT (**e**) (one-way ANOVA with Dunnett's test [**a**,**c**], two-way ANOVA [**b**,**e**]). Error bars represent the mean ± standard error of the mean (SEM). n.s. : not significant.

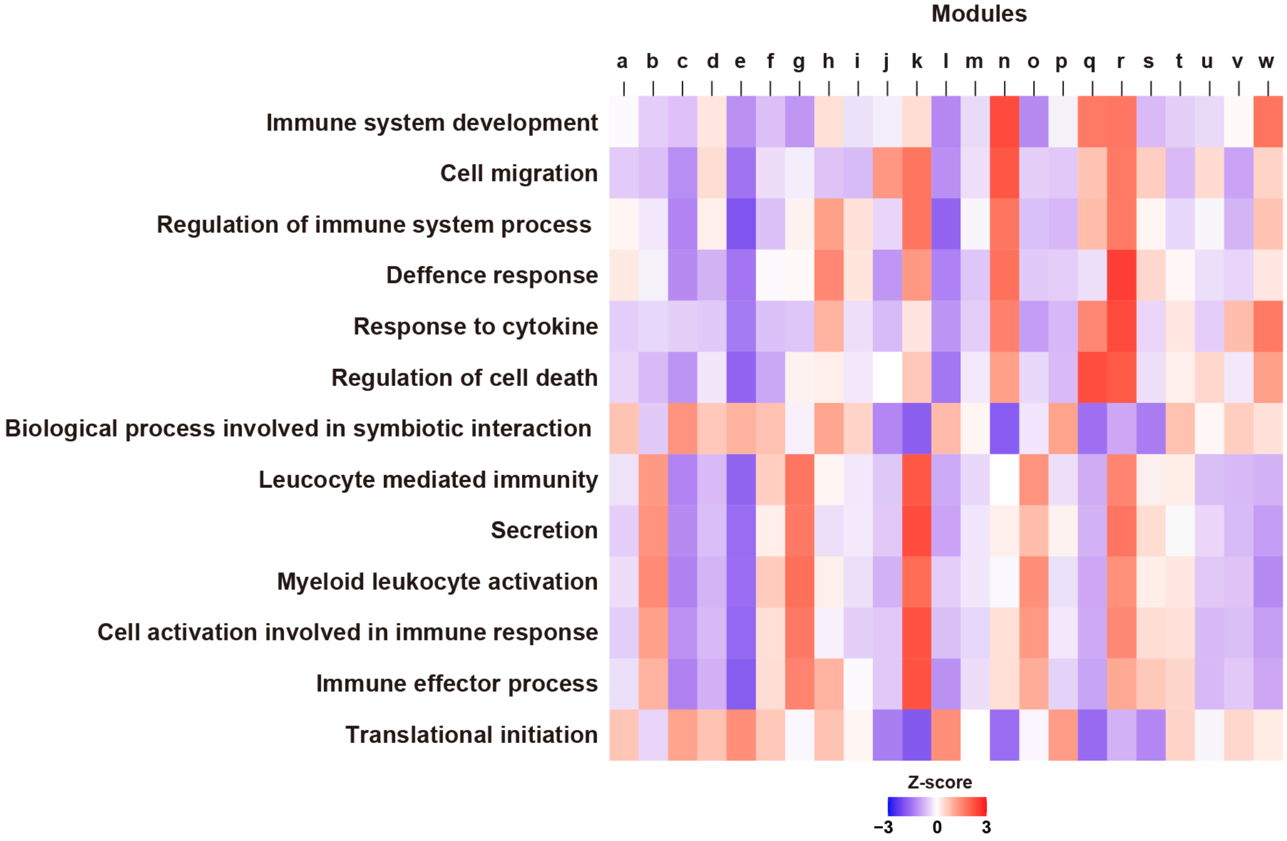


**Supplementary Figure 2.**

Enrichment of gene ontologies in each module illustrating cellular characteristics of myeloid cells (shown in **Fig.3f**) in WT and *App*$^{NL\text{-}G\text{-}F/NL\text{-}G\text{-}F}$ mice with/without ischemic stroke. Heatmap shows the Z-score of GSEA enrichment score: a higher Z-score means more enrichment of indicated gene ontologies.

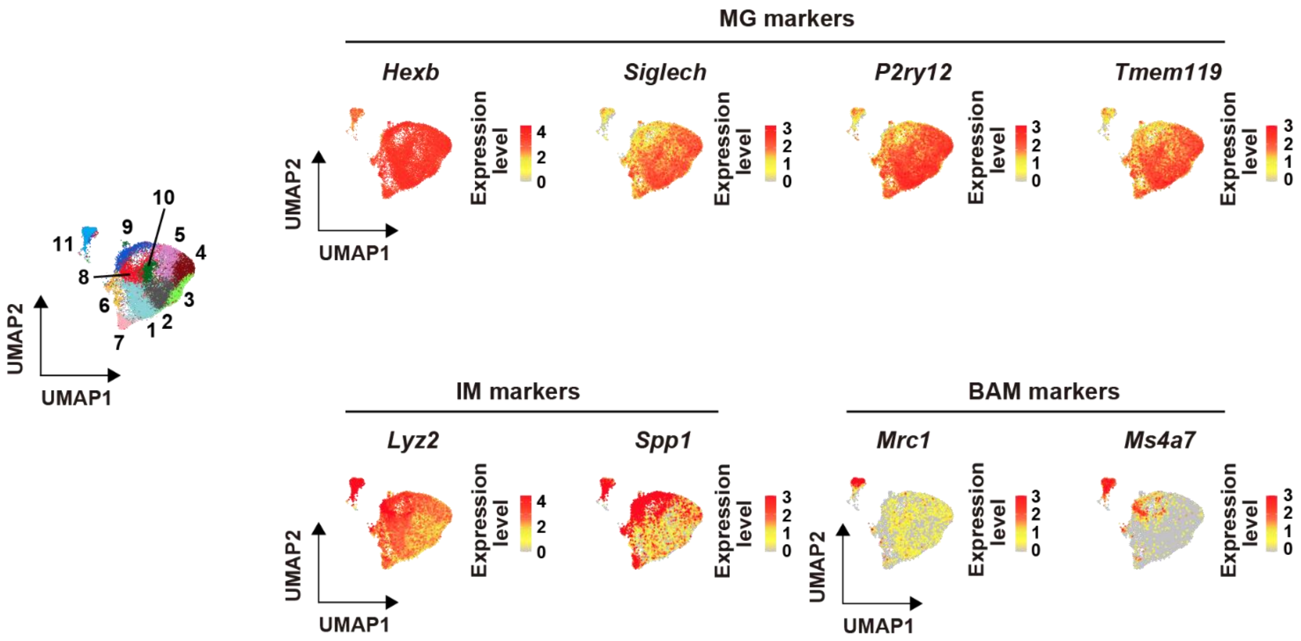


**Supplementary Figure 3.**

The nature of myeloid cells classified by marker gene expression of microglia (MG), infiltrating myeloid cell (IM), and border-associated macrophage (BAM). The left panel shows the integrated UMAP of myeloid cells in WT and *App*$^{NL\text{-}G\text{-}F/NL\text{-}G\text{-}F}$ mice with/without ischemic stroke. Right panel shows the gene expression levels of each myeloid cell marker (previously described elsewhere[1,2]).

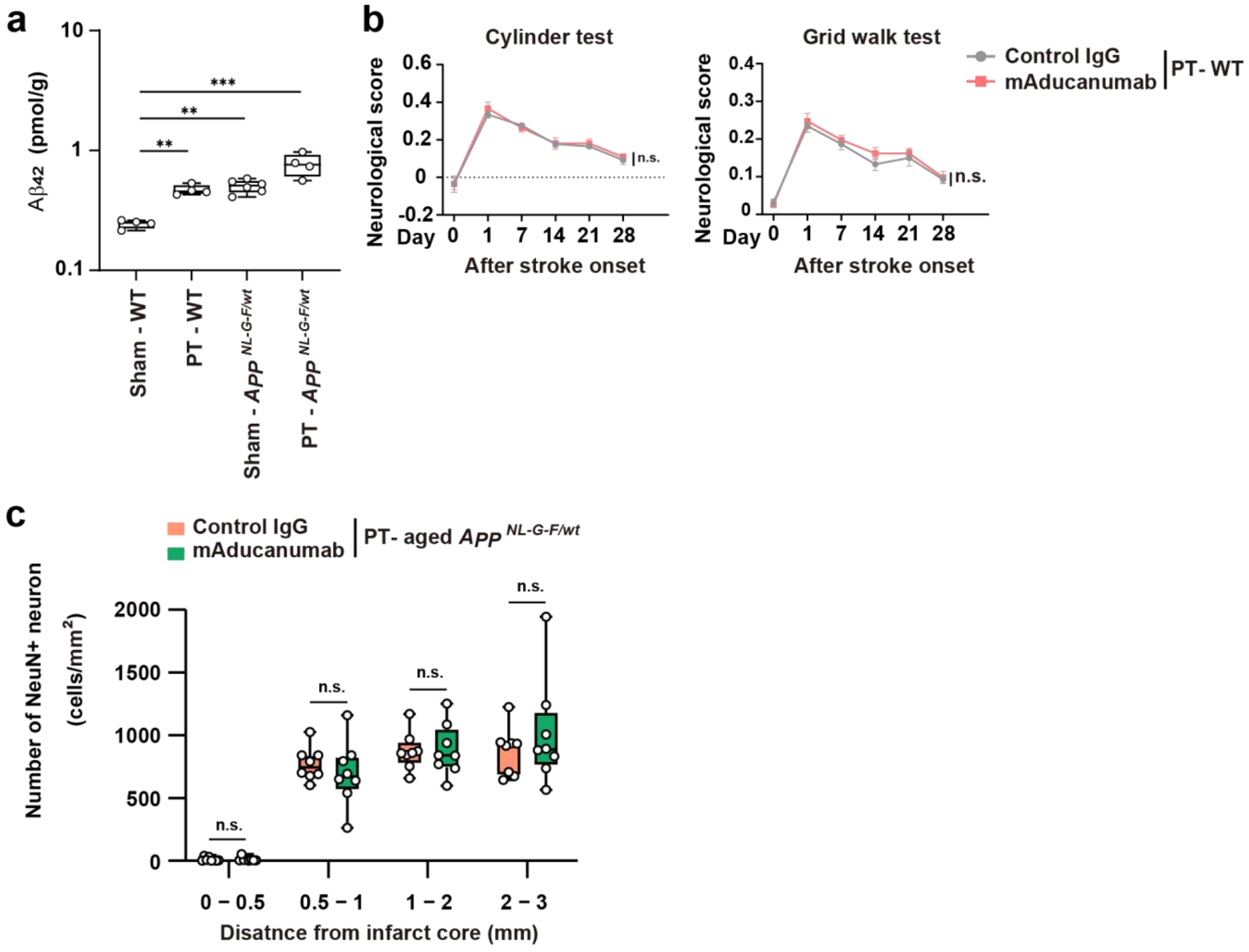


**Supplementary Figure 4.**

(**a**) Quantification of A$\beta_{42}$ peptide [pmol] in the brain tissue [g] by ELISA. The antibody used in the ELISA kit cross-reacted between mouse and human A$\beta_{42}$ peptide. (**b**) The comparison of neurological deficits between WT mice administered control IgG and mAducanumab 24 hours after ischemic stroke (n = 5 for control IgG, n = 8 for mAducanumab). (**c**) The number of NeuN$^+$ neurons in each cortical region away from the ischemic core as specified in **Fig.4h**. $^{**}p < 0.01$, $^{***}p < 0.001$ vs. sham-WT (**a**) (one-way ANOVA with Dunnett's test [**a**], two-way ANOVA [**b**], two-way ANOVA with Sidak's multiple comparison test [**c**]). Error bars represent the mean ± standard error of the mean (SEM). n.s. : not significant.

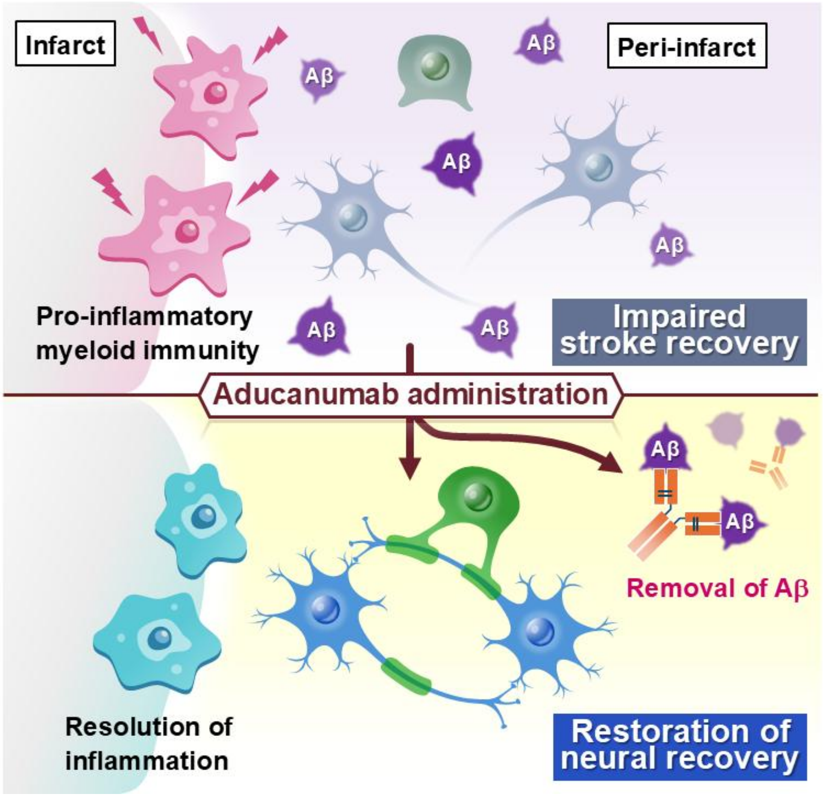


**Supplementary Figure 5.**

Unique pro-inflammatory myeloid immunity in the ischemic brain with Aβ accumulation impaired the neural gene expression associated with recovery processes after ischemic stroke. Post-stroke administration of anti-Aβ antibody resolved such inflammation and restored neural repair after ischemic stroke.

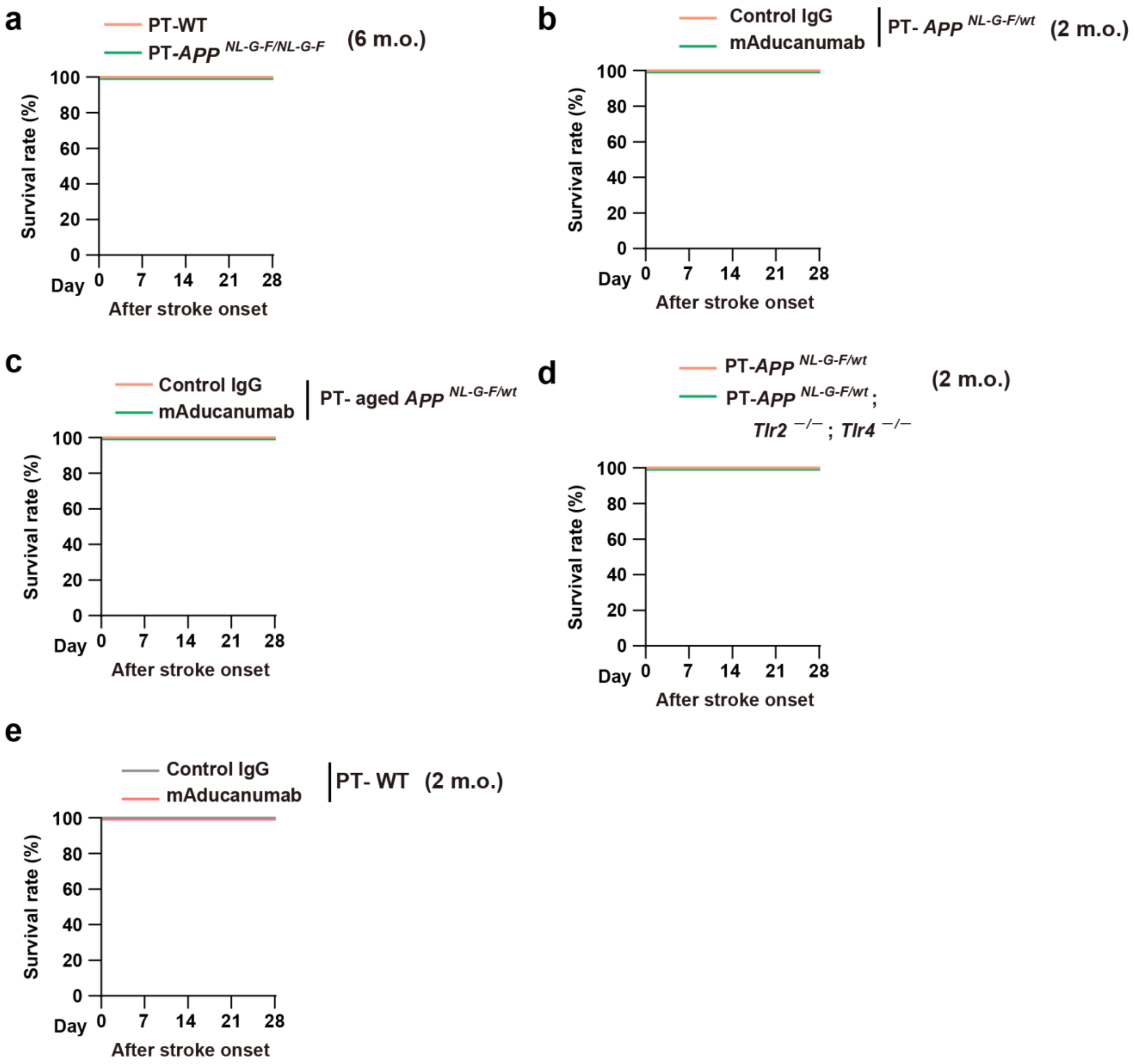


**Supplementary Figure 6.**

(**a**) The survival rate of six-month-old WT, *App*$^{NL\text{-}G\text{-}F/wt}$ , and *App*$^{NL\text{-}G\text{-}F/NL\text{-}G\text{-}F}$ mice after ischemic stroke onset. (**b**,**c**) The survival rate compared between the administration of control IgG and mAducanumab 24 hours after ischemic stroke in two-month-old *App*$^{NL\text{-}G\text{-}F/wt}$ mice (**b**), or in aged *App*$^{NL\text{-}G\text{-}F/wt}$ mice (**c**). (**d**) The survival rate of two-month-old *App*$^{NL\text{-}G\text{-}F/wt}$ mice and *Tlr2*$^{-/-}$; *Tlr4*$^{-/-}$; *App*$^{NL\text{-}G\text{-}F/wt}$ mice after ischemic stroke onset. (**e**) The comparison of survival ratio between the administration of control IgG and mAducanumab 24 hours after ischemic stroke in two-month-old WT mice.

| | WT | $App^{NL\text{-}G\text{-}F/NL\text{-}G\text{-}F}$ |
|---|---|---|
| MABP (mmHg) | 82 ± 3 | 81 ± 3 |
| pH | 7.41 ± 0.02 | 7.38 ± 0.02 |
| $PaO_2$ (mmHg) | 144 ± 15 | 135 ± 5 |
| $PaCO_2$ (mmHg) | 23.8 ± 1.4 | 27.8 ± 2.1 |
| Hematocrit (%) | 40 ± 2 | 37 ± 1 |
| Glucose (mg/dl) | 206 ± 6 | 211 ± 14 |

**Supplementary Table 1.**

Physiological data of WT and $App^{NL\text{-}G\text{-}F/NL\text{-}G\text{-}F}$ mice.

| No. | Age/sex | Etiology, region of brain infarction | Cause of death | Comorbidities | tPA administration |
|---|---|---|---|---|---|
| 1 | 70 Female | Embolic, left MCA territory | Respiratory arrest due to cerebral herniation | HCV hepatitis | Not performed |
| 2 | 80 Female | Atherothrombotic, left MCA territory | Rupture of aortic dissecting aneurysm | Congestive heart failure<br>Hypertension<br>Arrhythmia (VPC) | Not performed |
| 3 | 63 Male | Cardioembolic, right MCA territory | Respiratory arrest due to cerebral herniation | Atrial fibrillation<br>Diabetes mellitus<br>Chronic kidney disease | Not performed |
| 4 | 77 Male | Embolic, bilateral thalamus, multiple | Respiratory arrest | Atrial fibrillation | Not performed |
| 5 | 68 Female | Cardioembolic, left MCA, PCA territory | Respiratory arrest due to cerebral herniation | Rheumatoid arthritis<br>Congestive heart failure | Not performed |
| 6 | 62 Male | Atherothrombotic, right MCA territory | Respiratory arrest due to cerebral herniation | Hypertension<br>Congestive heart failure<br>Chronic kidney disease<br>Severe cognitive deficit | Not performed |
| 7 | 69 Male | Cardioembolic, right MCA territory | Respiratory arrest due to cerebral herniation | Sick sinus syndrome | Not performed |
| 8 | 74 Female | Cardioembolic, right ACA, MCA territory | Respiratory arrest due to cerebral herniation | Atrial fibrillation | Not performed |
| 9 | 80 Female | Cardioembolic, right MCA territory | Respiratory arrest due to cerebral herniation | Atrial fibrillation<br>Hypertension<br>Mild cognitive deficit | Not performed |
| 10 | 69 Female | Embolic, left MCA territory | Respiratory arrest | Hepatocellular carcinoma<br>Behcet's disease | Not performed |
| 11 | 81 Female | Cardioembolic, left MCA territory | Aspiration pneumonia | Atrial fibrillation<br>Hypertension | Not performed |

Abbreviations:
ACA: anterior cerebral artery, MCA: middle cerebral artery, PCA: posterior cerebral artery, ICA: internal carotid artery

### Supplementary Table 2.

Detailed clinical information on ischemic stroke patients (related to Figure 2).
